# Excess PrP^C^ inhibits muscle cell differentiation via miRNA-enhanced liquid–liquid phase separation implicated in myopathy

**DOI:** 10.1101/2023.02.06.527278

**Authors:** Jing Tao, Yanping Zeng, Bin Dai, Yin Liu, Xiaohan Pan, Li-Qiang Wang, Jie Chen, Yu Zhou, Zuneng Lu, Liwei Xie, Yi Liang

## Abstract

The cellular prion protein (PrP^C^), a glycoprotein existing in membrane-bound and cytoplasmic forms, has functional importance in skeletal muscle, but the mechanism behind the phenomenon remains poorly understood. Here we report that PrP^C^ is overexpressed and located in the cytoplasm of the skeletal muscle of six myopathy patients; cytoplasmic PrP^C^ strongly inhibits skeletal muscle cell autophagy and blocks myoblast differentiation. PrP^C^ selectively binds to a subset of miRNAs during myoblast differentiation, and the co-localization of PrP^C^ with miR-214-3p was clearly observed in the skeletal muscle of six myopathy patients but not in that of four age-matched controls. We demonstrate that PrP^C^ is overexpressed in skeletal muscle cells under pathological conditions and inhibits muscle cell differentiation via physically interacting with a subset of miRNAs to significantly inhibit autophagy-related protein 5-dependent autophagy, and selectively recruits these miRNAs into phase-separated condensates in living myoblasts, which in turn greatly enhances liquid–liquid phase separation (LLPS) of PrP^C^ and results in the subsequent PrP aggregation and muscle bundle formation in myopathy patients characterized by incomplete muscle regeneration. Our findings show how excess PrP^C^ can inhibit muscle cell differentiation via miRNA-enhanced LLPS implicated in myopathy.

---

Mammalian prion protein (PrP) has two forms distinct in their structure and function, the cellular prion protein (PrP^C^) and its pathological aggregated form PrP^Sc^ (refs. ^1−6^). Prion diseases are a group of transmissible neurodegenerative diseases primarily caused by the conformational conversion of PrP from α-helix-dominant PrP^C^ to β-sheet−rich PrP^Sc^ in many mammal species (refs. ^1−12^). The benign cellular form PrP^C^, a glycosylphosphatidylinositol (GPI)-anchored glycoprotein existing in membrane-bound and cytoplasmic forms with multifaceted functions^13,14^, has functional importance in skeletal muscle^15−22^, such as maintaining normal muscle size and function^15^, promoting muscle regeneration^16^, and linking to myoblast differentiation^15,18^, and PrP^C^ dysfunction is observed in the skeletal muscle of patients with inclusion-body myositis, dermatomyositis, and other myopathies^19,20,22^ or transgenic mice developing a rapidly progressive primary myopathy^21^, but the mechanism behind the phenomenon is still unclear.

Autophagy, an evolutionary conservative catabolic pathway through lysosomal degradation of intracellular components^23,24^, appears to increase and is required during myoblast differentiation^25,26^ and related to muscle pathophysiology^27^. Importantly, impaired autophagy is observed in aged muscle satellite cells^28^. Moreover, scientists have identified at least 35 genes coding autophagy-related (ATG) proteins, including autophagy-related protein 5 (ATG5) and microtubule-associated protein 1 light chain 3 (LC3), among which the E3 ubiquitin ligase ATG5 is essential for autophagosome elongation^23,24,29−32^. Currently, the pathological prion aggregates have been shown to trigger autophagy in skeletal muscle and can be degraded by autophagy^33^. However, it is unclear whether the benign cellular form PrP^C^ regulates autophagy and differentiation of skeletal muscle cells.

MicroRNAs (miRNAs) are small non-coding RNAs that are highly enriched in skeletal muscle and highly conserved from plants to mammals^34,35^. MiRNAs play roles in regulating differentiation, atrophy, and regeneration of skeletal muscle via interaction with specific proteins^35−38^. Three miRNAs, miR-181a-5p, miR-324-5p, and miR-451a, are overexpressed in the skeletal muscle of patients with spinal muscular atrophy, suggesting that targeting the overexpressed miRNAs may be a novel therapeutic approach against the disease^37^. However, it is unclear whether miRNAs regulate autophagy and differentiation of skeletal muscle cells via specific interaction with PrP^C^.

Protein and RNA molecules tend to form supramolecular assemblies called membrane-less organelles via liquid–liquid phase separation (LLPS) of proteins in cells to perform key functions^39−43^. The liquid droplets formed by biological macromolecules, called biomolecular condensates, have fusion properties^40,44^ and RNA may control or buffer LLPS of proteins with natively unfolded and/or low-complexity domains^43−46^. Because PrP^C^ contains a low-complexity, intrinsically disordered region (IDR) in its N-terminal domain, it undergoes LLPS in vitro and forms protein condensates^47−56^. PrP^C^ liquid-phase condensation is modulated by amyloid-β oligomers, neutralizing mutations, pathological mutations, RNA (polyU RNA, crude tRNA, and yeast total RNA), and other factors^47−56^. However, it is unclear whether PrP^C^ undergoes LLPS in vivo and whether the LLPS of PrP^C^ leads to pathological aggregation of the protein in cells. It also remains unknown whether PrP^C^ selectively recruits specific miRNAs into phase-separated condensates, which in turn regulates the LLPS of PrP^C^.

Here we report that PrP^C^ is overexpressed and located in the cytoplasm of the skeletal muscle of myopathy patients. We demonstrate that PrP^C^ located in the cytoplasm blocks muscle cell differentiation via selectively recruiting a subset of miRNAs including miR-214-3p into phase-separated condensates in living myoblasts, which in turn greatly enhances the LLPS of PrP^C^ and the subsequent PrP aggregation in vivo and significantly inhibits ATG5-dependent autophagy. Our findings provide insights into the regulation of PrP^C^ on muscle cell differentiation under pathological conditions via LLPS of the protein enhanced by a subset of miRNAs such as miR-214-3p, which has important implications in myopathy etiology.

## Results

### PrP^C^ overexpression is observed in the skeletal muscle of myopathy patients

We first took confocal images of frozen skeletal muscle sections from six myopathy patients and four age-matched controls (Fig. 1a,b). These myopathy patients include two patients with dermatomyositis, two patients with neurogenic myopathy, and two patients with muscular dystrophy, and those controls include one healthy individual, two patients with lipid storage myopathy, and one patient with glycogen storage disease (Table 1 and Fig. 1a−d). The frozen skeletal muscle sections were immunostained with the anti-PrP antibody 8H4 (red), stained with DAPI (blue), and visualized by confocal microscopy. PrP^C^ (red) was overexpressed and mainly located in the cytoplasm of the skeletal muscle of these myopathy patients (Fig. 1a). In sharp contrast, PrP^C^ overexpression was not observed in the skeletal muscle of those controls (Fig. 1b). H&E staining of the frozen skeletal muscle sections revealed that the skeletal muscle of these myopathy patients was characterized by incomplete muscle regeneration (cases 2 and 6 with obvious muscular atrophy) but muscle regeneration was not observed in the skeletal muscle of those controls (Fig. 1c,d). Interestingly, increased PrP^C^ expression is seen in inclusion-body myositis, dermatomyositis, and other myopathies^19−22^, in agreement with our experimental observations (Fig. 1a).

**Fig. 1.**
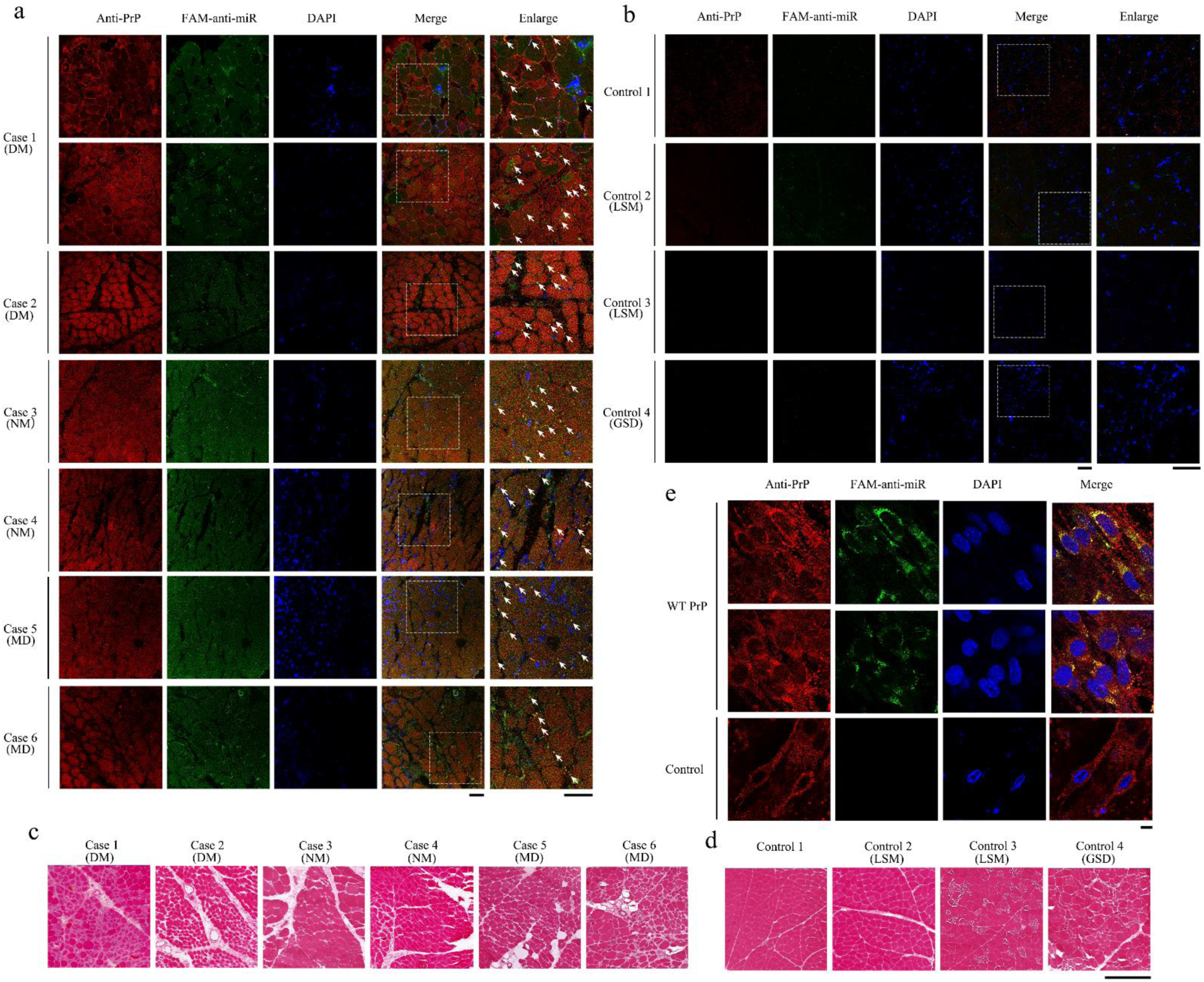
PrP^C^ overexpression and the co-localization of PrP^C^ with miR-214-3p were clearly observed in the skeletal muscle of myopathy patients but not in that of age-matched controls. **a**,**b**, Confocal images of frozen skeletal muscle sections from six myopathy patients (**a**) and four age-matched controls (**b**). These myopathy patients include two patients with dermatomyositis (DM) (Cases 1 and 2), two with neurogenic myopathy (NM) (Cases 3 and 4), and two with muscular dystrophy (MD) (Cases 5 and 6). The following are the controls: one healthy individual (Control 1), two patients with lipid storage myopathy (LSM) (Controls 2 and 3), and one with glycogen storage disease (GSD) (Control 4). The frozen skeletal muscle sections, in which miR-214-3p was detected by FISH (green) using an FAM-labeled miR-214-3p probe (FAM-anti-miR), were immunostained with the anti-PrP antibody 8H4 (red), stained with DAPI (blue), and visualized by confocal microscopy. The enlarged regions (right) show 4-fold enlarged images from the merged images. Scale bars, 75 μm. **a**, Excess PrP^C^ (red) mainly located in the cytoplasm, abundant miR-214-3p (green dots), and the co-localization of excess PrP^C^ with miR-214-3p (yellow dots in the merged images highlighted by using white arrows) were observed in the skeletal muscle of these myopathy patients. **b**, PrP^C^ overexpression, miR-214-3p expression, and the co-localization of PrP^C^ with miR-214-3p (the merged images) were all not observed in the skeletal muscle of those controls. **c**,**d**, H&E staining of the frozen skeletal muscle sections revealed that the skeletal muscle of these myopathy patients was characterized by incomplete muscle regeneration (cases 2 and 6 with obvious muscular atrophy) (**c**) but muscle regeneration was not observed in the skeletal muscle of those controls (**d**). Scale bar, 400 μm. **e**, C2C12 mouse myoblasts (control) and C2C12 myoblasts stably expressing WT PrP^C^ upon differentiation for 4 days, in which miR-214-3p was detected by FISH (green) using FAM-anti-miR, were immunostained with the anti-PrP antibody 8H4 (red), stained with DAPI (blue), and visualized by confocal microscopy. Excess PrP^C^ (red) mainly located in the cytoplasm, abundant miR-214-3p (green dots), and the co-localization of excess PrP^C^ with miR-214-3p (yellow dots in the merged images) were observed in differentiating C2C12 cells stably expressing WT PrP^C^. Both miR-214-3p expression and the co-localization of endogenous PrP^C^ with miR-214-3p (the merged images), however, were not observed in differentiating C2C12 cells (control). Scale bar, 10 μm.

**Table 1.** Clinical details of patients with dermatomyositis, neurogenic myopathy, and muscular dystrophy at the time when skeletal muscle samples were taken.

| Case | Gender | Age | Mean age ± S.D. | Clinical diagnosis |
| --- | --- | --- | --- | --- |
| Myopathy |  |  |  |  |
| 1 | F | 52 | 50.9 ± 4.9 | Dermatomyositis |
| 2 | F | 49 |  | Dermatomyositis |
| 3 | M | 51 |  | Neurogenic myopathy |
| 4 | F | 58 |  | Neurogenic myopathy |
| 5 | M | 43 |  | Muscular dystrophy |
| 6 | F | 49 |  | Muscular dystrophy |
| Control |  |  |  |  |
| 1 | F | 70 | 57.3 ± 17.1 | Healthy individual |
| 2 | M | 32 |  | Lipid storage myopathy |
| 3 | M | 64 |  | Lipid storage myopathy |
| 4 | F | 63 |  | Glycogen storage disease |

### Maintenance of PrP^C^ homeostasis is essential for myoblast differentiation

A central hypothesis for PrP^C^’s function is that its overexpression or knockout may impair its function in regulating myoblast differentiation. We have shown that PrP^C^ overexpression does impair its function in regulating myoblast differentiation (Fig. 1a-d). To test this hypothesis at the cellular level, we used murine-derived C2C12 myoblast cells, which provide a fascinating possibility as myoblasts are proliferative but differentiate into multinucleated, elongated myotubes after serum deprivation^57^. Moreover, PrP^C^ expression is upregulated during myoblast differentiation^58^. C2C12 myoblasts stably overexpressing full-length wild-type (WT) mouse PrP^C^ (Fig. 2b,e,f), C2C12 myoblasts stably overexpressing F198S PrP^C^ (Fig. 2c,e,f), a genetic Gerstmann−Sträussler−Scheinker disease−related mutation^7,12^, and C2C12 myoblasts knockout (KO) for PrP^C^ (Fig. 2d−f) were cultured until confluence was reached 90% and then incubated with a differentiation medium for up to 6 days (Fig. 2a−d) or 3 and 5 days, respectively (Fig. 2e,f), using C2C12 myoblasts as a control (Fig. 2a,e,f). Here we used mouse PrP^C^ instead of the human counterpart because these two proteins have more than 90% sequence identity^59^. We used anti-MyHC antibody and anti-MyoG antibody to detect the expression of myogenic differentiation markers, myosin heavy chain (MyHC) and myogenin (MyoG), respectively. The cell lysates were probed by the anti-PrP monoclonal antibody 8H4, anti-MyHC antibody, anti-MyoG antibody, and anti-β-actin antibody, respectively (Fig. 2a−d). The above cells were immunostained with the anti-PrP antibody 8H4 (red) (Fig. 2e) or the anti-MyHC antibody MF20 (red) (Fig. 2f), stained with DAPI (blue), and observed by confocal microscopy. Endogenous PrP^C^ located on the plasma membrane, including its diglycosylated, monoglycosylated, and unglycosylated forms (Fig. 2a,e), and myoblast differentiation (Fig. 2a,f) were both observed when C2C12 cells (control) were incubated with the differentiation medium for 3 days. After 5 days of differentiation, however, endogenous PrP^C^ located on the plasma membrane, endogenous PrP^C^ located in the cytoplasm (Fig. 2e), and abundant multinucleated, elongated myotubes (Fig. 2f) differentiated from C2C12 myoblasts were observed. Importantly, excess WT PrP^C^ (Fig. 2b,e,f) and excess familial mutation F198S PrP^C^ (Fig. 2c,e,f) located in the cytoplasm (Fig. 2e) both strongly inhibited and blocked muscle cell differentiation when C2C12 myoblasts stably expressing PrP^C^ were incubated with the differentiation medium for 3, 4, 5, and 6 days. After 3 days of differentiation, excess WT PrP^C^ is partly located on the plasma membrane and partly located in the cytoplasm. After 5 days of differentiation, however, excess WT PrP^C^ is completely located in the cytoplasm (Fig. 2e). Moreover, PrP^C^ deficiency completely blocked muscle cell differentiation when C2C12 myoblasts KO for PrP^C^ were incubated with this medium for 3, 4, 5, and 6 days (Fig. 2d,e,f). Together, the data showed that maintenance of PrP^C^ homeostasis is essential for myoblast differentiation.

**Fig. 2.**
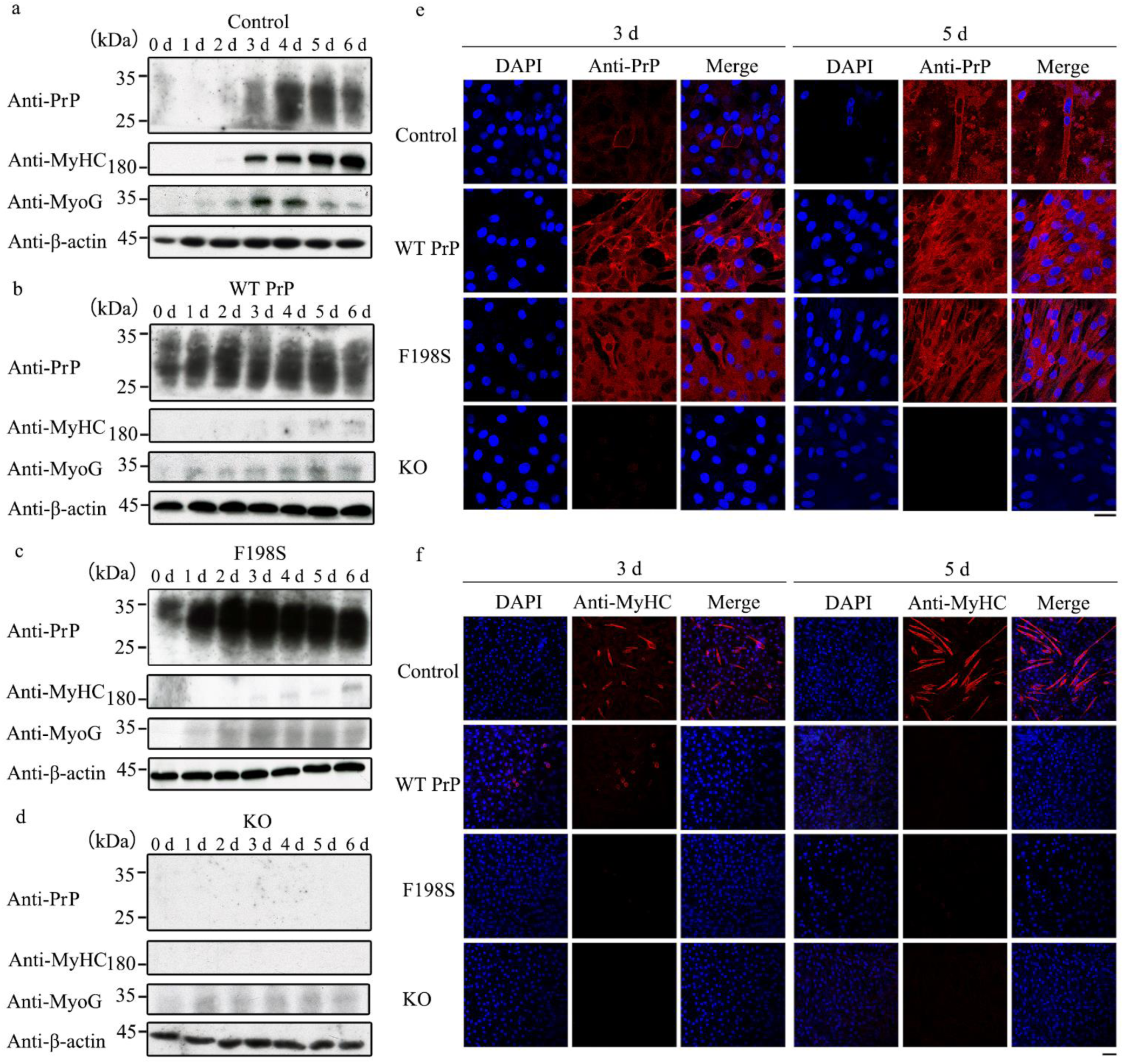
Maintenance of PrP^C^ homeostasis is essential for myoblast differentiation. **a**–**d**, C2C12 mouse myoblasts (control) (**a**), C2C12 myoblasts stably expressing full-length wild-type mouse PrP^C^ (WT PrP^C^) (**b**), C2C12 myoblasts stably expressing F198S PrP^C^ (**c**), and C2C12 myoblasts KO for PrP^C^ (**d**) were cultured until confluence was reached 90% and then incubated with a differentiation medium supplemented with 2% (v/v) horse serum (Gibco) and 1% (v/v) penicillin-streptomycin for 0, 1, 2, 3, 4, 5, and 6 days, respectively. The cell lysates from the above cells were probed by the anti-PrP monoclonal antibody 8H4, anti-MyHC antibody, anti-MyoG antibody, and anti-β-actin antibody, respectively. All blots also show the position of the molecular-weight markers. **e**,**f**, The above four cell lines were incubated with the differentiation medium for 3 and 5 days, respectively, then immunostained with the anti-PrP antibody 8H4 (red) (e) or the anti-MyHC antibody MF20 (red) (**f**), stained with DAPI (blue), and observed by confocal microscopy. Scale bars, 50 (**e**) and 75 (**f**) μm, respectively. **a**, Both endogenous PrP^C^ and myoblast differentiation were observed until C2C12 cells (control) were incubated with the differentiation medium for 3 days. Endogenous PrP^C^ located on the plasma membrane (**e**) and myoblast differentiation (**f**) were both observed when C2C12 cells (control) were incubated with the differentiation medium for 3 days. After 5 days of differentiation, however, endogenous PrP^C^ located on the plasma membrane, endogenous PrP^C^ located in the cytoplasm (**e**), and abundant multinucleated, elongated myotubes (**f**) differentiated from C2C12 myoblasts were observed. Importantly, excess WT PrP^C^ (**b**,**e**,**f**) and excess familial mutation F198S PrP^C^ (**c**,**e**,**f**) located in the cytoplasm (**e**) both strongly inhibit and block muscle cell differentiation when C2C12 myoblasts stably expressing PrP^C^ were incubated with the differentiation medium for 3, 4, 5, and 6 days. **d**,**e**,**f**, PrP^C^ deficiency completely blocks muscle cell differentiation when C2C12 myoblasts KO for PrP^C^ were incubated with this medium for 3, 4, 5, and 6 days. The F198S mutation in human corresponds to the F197S mutation in mouse.

### Excess PrP^C^ strongly inhibits skeletal muscle cell autophagy and blocks myoblast differentiation

Given that the benign cellular form PrP^C^ does precisely regulate the differentiation of skeletal muscle cells (Fig. 2), we predicted that PrP^C^ might directly regulate skeletal muscle cell autophagy, an important pathway for cell differentiation. We next used confocal microscopy and Western blotting to test this hypothesis. The co-localization of endogenous PrP^C^, which is mainly attached to the plasma membrane, with the autophagy marker LC3B (yellow dots in the merged image, Fig. 3a) was observed in C2C12 cells (control) upon differentiation for 3 days. The co-localization of endogenous PrP with LC3B (yellow dots in the merged image, Fig. 3b) was also observed in the elongated myotubes differentiated from C2C12 myoblasts after 5 days of differentiation. After 3 days of differentiation, endogenous PrP^C^ did form abundant LC3B-positive puncta (green dots, Fig. 3a) in C2C12 cells, and after 5 days of differentiation, endogenous PrP^C^ produced more abundant LC3B-positive puncta (green dots, Fig. 3b) in C2C12 cells, as detected by immunofluorescence using anti-LC3B antibody (green) and the anti-PrP antibody 8H4 (red). Importantly, excess PrP^C^ strongly inhibited skeletal muscle cell autophagy and blocked myoblast differentiation, producing much fewer LC3B-positive puncta (green dots, Fig. 3a,b) when C2C12 cells stably expressing WT PrP^C^ were incubated with the differentiation medium for 3 and 5 days. After 3 days of differentiation, excess PrP^C^ is partly located on the plasma membrane and partly located in the cytoplasm (Fig. 3a); after 5 days of differentiation, however, excess PrP^C^ is completely located in the cytoplasm (Fig. 3b); under both conditions, the co-localization of PrP^C^ and LC3B (the merged images, Fig. 3a,b) were not observed. Moreover, PrP^C^ deficiency completely blocked muscle cell differentiation but did not inhibit skeletal muscle cell autophagy (green dots, Fig. 3a,b) when C2C12 myoblasts KO for PrP^C^ were incubated with this medium for 3 and 5 days. To gain a quantitative understanding of how PrP^C^ regulates skeletal muscle cell autophagy, we detected two autophagy-related proteins ATG5 and LC3B from the above cells using anti-ATG5 and anti-LC3B antibodies, respectively (Fig. 3c). Upon differentiation for 3 and 5 days, the relative amounts of ATG5 and LC3B-II in the cell lysates from C2C12 myoblasts stably expressing WT PrP^C^ were significantly lower than those in the control cell lysates from C2C12 myoblasts (orange) (*p* < 0.05) (Fig. 3d,e). Thus, excess PrP^C^ significantly inhibited ATG5-dependent and LC3B-dependent autophagy in differentiating C2C12 cells stably expressing WT PrP^C^, compared to those in differentiating C2C12 myoblasts (control). Moreover, PrP^C^ deficiency increased skeletal muscle cell autophagy, when C2C12 myoblasts KO for PrP^C^ were incubated with this medium for 3 and 5 days, respectively (Fig. 3a,b,d,e,). Intriguingly, we observed that the levels of ATG5 and LC3B-II were strongly enhanced during myoblast differentiation (Fig. 3d,e). These results demonstrate that excess PrP^C^ strongly inhibits skeletal muscle cell autophagy and blocks myoblast differentiation and indicate that PrP^C^ deficiency increases skeletal muscle cell autophagy but completely blocks muscle differentiation.

**Fig. 3.**
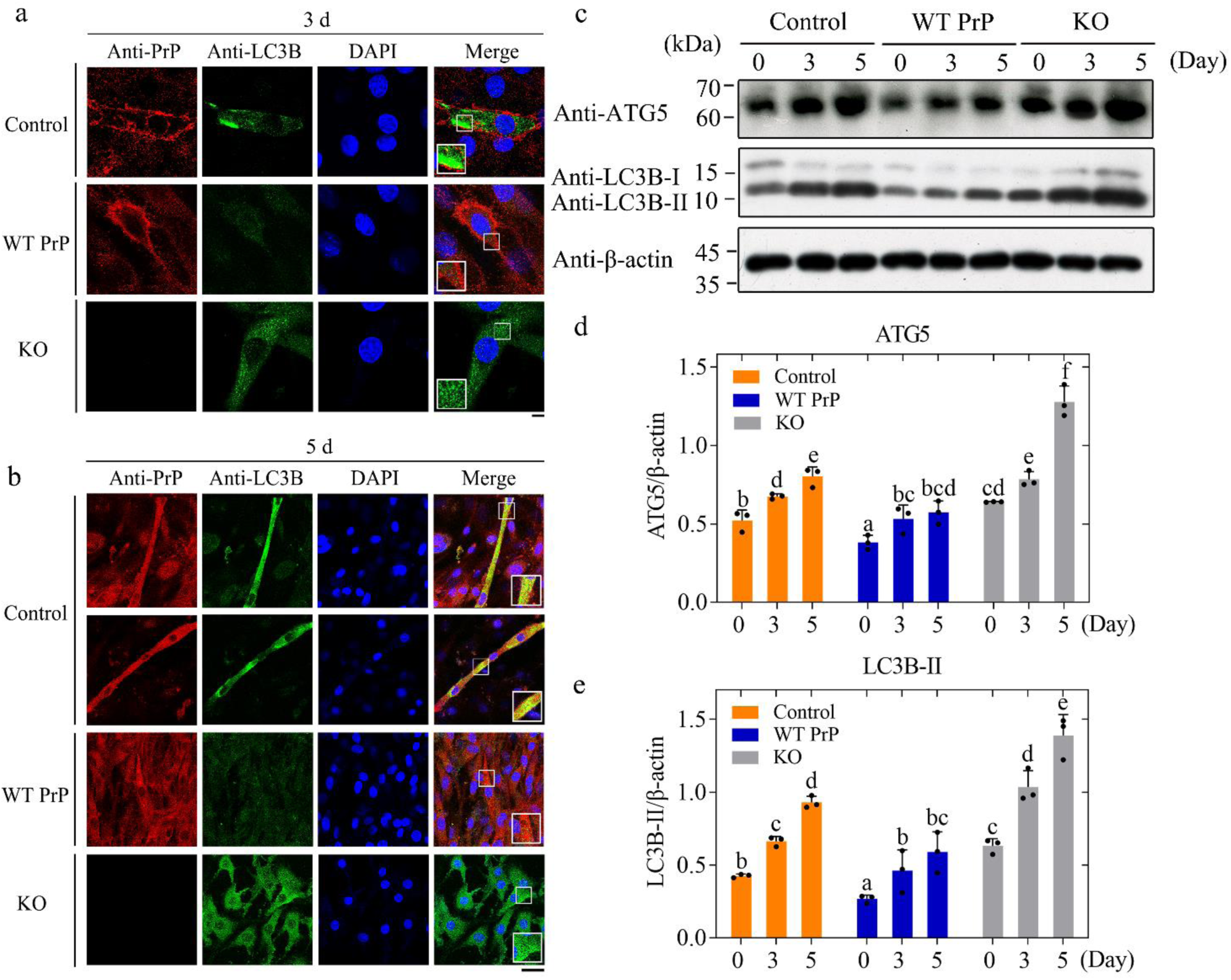
Excess PrP^C^ strongly inhibits skeletal muscle cell autophagy and blocks myoblast differentiation. **a**,**b**, C2C12 mouse myoblasts (control), C2C12 myoblasts stably overexpressing WT PrP^C^, and C2C12 myoblasts KO for PrP^C^ were cultured until confluence was reached 90% and then incubated with the differentiation medium for 3 (**a**) and 5 (**b**) days, respectively. C2C12 cells were coimmunostained with anti-LC3B antibody (green) and the anti-PrP antibody 8H4 (red), stained with DAPI (blue), and visualized by confocal microscopy. The enlarged regions in the lower left corner (**a**) or the lower right corner (**b**) of the merged images show 4-fold enlarged images from the same images. Scale bars, 7.5 (**a**) and 50 (**b**) μm, respectively. **a**, Endogenous PrP^C^ mainly attached to the plasma membrane, abundant LC3B-positive puncta (green dots), and the co-localization of endogenous PrP^C^ with the autophagy marker LC3B (yellow dots in the merged image) were observed when C2C12 cells (control) were incubated with the differentiation medium for 3 days. **b**, Endogenous PrP^C^, more abundant LC3B-positive puncta (green dots), and the co-localization of endogenous PrP^C^ with LC3B (yellow dots in the merged images) were all observed in the elongated myotubes differentiated from C2C12 myoblasts (control) after 5 days of differentiation. **a**,**b**, Importantly, excess PrP^C^ strongly inhibits skeletal muscle cell autophagy and blocks myoblast differentiation, producing much fewer LC3B-positive puncta (green dots) when C2C12 cells stably expressing WT PrP^C^ were incubated with the differentiation medium for 3 and 5 days. PrP^C^ deficiency completely blocks muscle cell differentiation when C2C12 myoblasts KO for PrP^C^ were incubated with this medium for 3 and 5 days. **c**, The above three cell lines were incubated with the differentiation medium for 3 and 5 days, respectively, and the cell lysates were probed by the anti-ATG5 antibody, anti-LC3B antibody, and anti-β-actin antibody, respectively. All blots also show the position of the molecular-weight markers. **d**,**e**, The relative amount of ATG5 (**d**) or LC3B-II (**e**) in the above cell lines (solid black circles shown in scatter plots) was determined as a ratio of the density of ATG5 or LC3B-II bands over the density of β-actin band in cell lysates and expressed as the mean ± S.D. (with error bars) of values obtained in three independent experiments. Importantly, excess PrP^C^ significantly inhibits ATG5-dependent and LC3B-dependent autophagy in differentiating C2C12 cells stably expressing WT PrP^C^ (blue), compared to those in differentiating C2C12 myoblasts (control) (orange). **a**,**b**,**d**,**e**, PrP^C^ deficiency increases skeletal muscle cell autophagy, when C2C12 myoblasts KO for PrP^C^ were incubated with this medium for 3 and 5 days, respectively. One-way ANOVA and multiple comparisons were performed by SPSS 19.0 and different letters indicate significant differences at the level of *p* < 0.05.

### PrP^C^ selectively binds to a subset of miRNAs during myoblast differentiation

To explore the mechanism underlying a potential relationship between PrP^C^ and miRNAs during myoblast differentiation, we first pursued PrP^C^-associated miRNAs in differentiating C2C12 cells. To this end, we performed formaldehyde crosslinking and RNA immunoprecipitation (RIP) using anti-mouse antibody-binding beads incubated with the anti-PrP antibody 8H4 or mouse IgG to pull down a subset of PrP^C^-bound miRNAs from C2C12 cells stably expressing WT PrP^C^ and C2C12 myoblasts (control) differentiated for 4 days, and identified many selectively enriched miRNAs based on sorted ratios of RIP to input. Notably, a subset of PrP^C^-bound miRNAs (red) was significantly enriched in WT PrP^C^ RIP upon differentiation for 4 days, compared to control RIP and two inputs (Fig. 4a). Our Western blotting experiments indicated that the antibody-binding beads IPed endogenous PrP^C^ in C2C12 myoblasts after 4 days of differentiation (Fig. 4b). Displaying the small RNA-seq data in two volcano plots (Fig. 4c,d) and using a log_2_ fold-change cutoff of 0.5 and a *P*-value cutoff of 0.05, we identified 51 miRNAs bound by PrP^C^ in differentiating C2C12 cells stably expressing WT PrP^C^ versus 31 miRNAs bound by endogenous PrP^C^ in differentiating C2C12 cells (control) (Supplementary Table 1). From a total of 519 and 455 miRNAs that we identified, 51 and 31 were significantly enriched in differentiating C2C12 cells stably expressing WT PrP^C^ (Fig. 4d) and differentiating C2C12 cells (control) (Fig. 4c), respectively, and 468 and 424 were depleted, respectively. Ten overlaps were identified between the two subsets of miRNAs (Fig. 4e), and these miRNAs were further verified by RT−qPCR (Fig. 4f,g) and Gene Ontology (GO) analyses (Fig. 4h,i). Forty-one miRNAs bound by PrP^C^, such as miR-214-3p, miR-204-5p, miR-499-5p, and miR-92b-3p, which have inhibitory effects on cell differentiation and autophagy^60−63^, were identified for WT PrP RIP only, and 21 miRNAs bound by endogenous PrP^C^, such as miR-486a-5p, miR-181b-5p, and miR-206-3p, which have enhancing effects on cell differentiation and autophagy^64−66^, were for control RIP only (Fig. 4c,d and Supplementary Table 1). A subset of PrP^C^-bound miRNAs, including miR-214-3p, miR-204-5p, and miR-499-5p, and a subset of endogenous PrP^C^-bound miRNAs such as miR-486a-5p, miR-181b-5p, and miR-206-3p, but not U6 or any of the non-target miRNAs tested, were significantly enriched in differentiating C2C12 cells stably expressing WT PrP^C^ (Fig. 4g) and the differentiating control cells (Fig. 4f), respectively. We then identified the top six biological processes (BP) of GO enrichment in the control cells, including “Regulation of cellular process”, “Regulation of cell differentiation”, and “Cell differentiation” (Fig. 4h), and the top eight BPs of GO enrichment in C2C12 cells stably overexpressing WT PrP^C^, including “Regulation of cellular process”, “Cell differentiation”, “Autophagy”, and “Process utilizing autophagic mechanism” (Fig. 4i). These significantly enriched pathways are related to cell differentiation and autophagy and may reflect the regulatory roles of miRNAs enriched by PrP^C^.

**Fig. 4.**
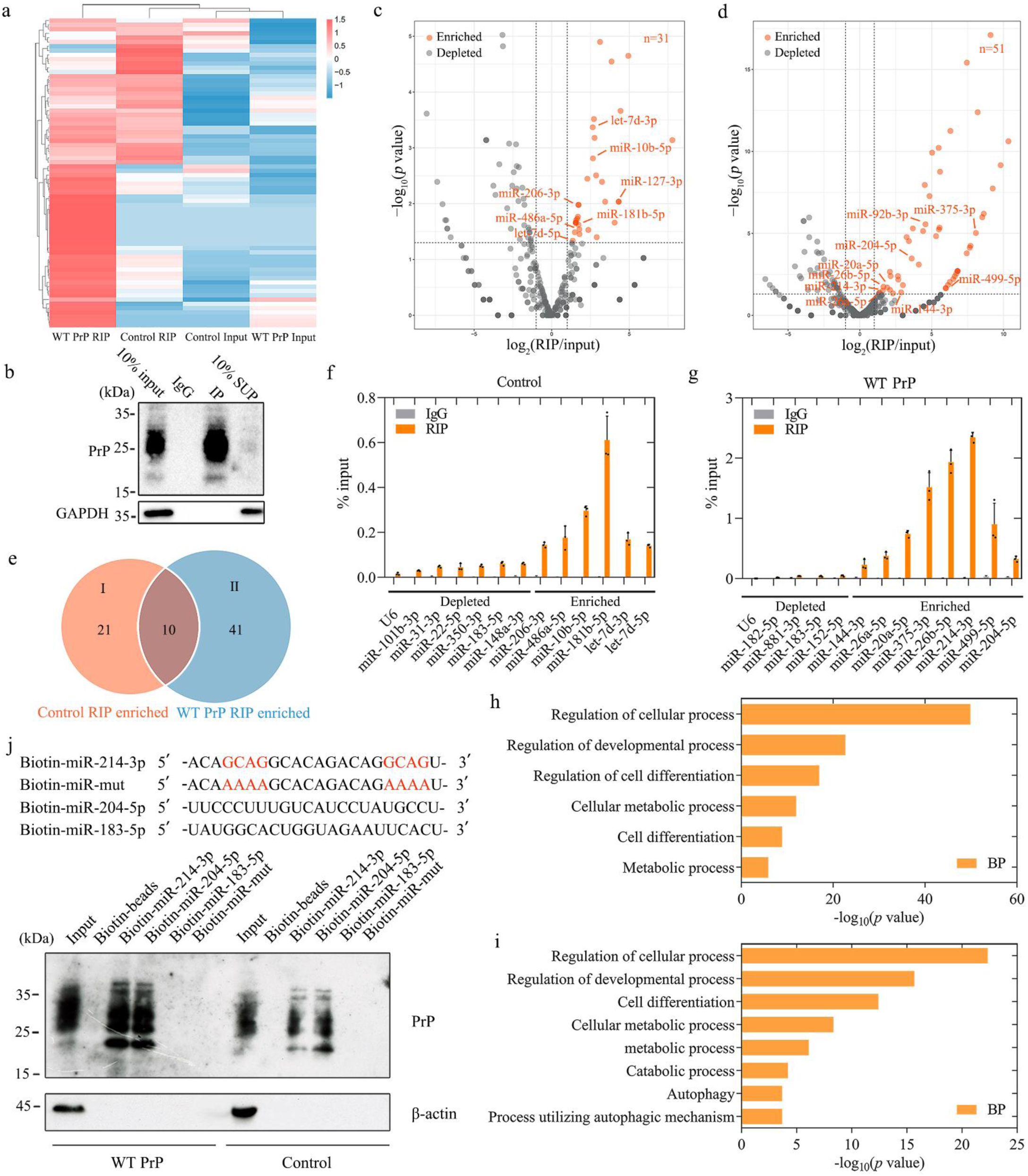
PrP^C^ selectively binds to a subset of miRNAs during myoblast differentiation. **a**, Profile of total miRNAs versus those enriched with PrP^C^ RIP. The anti-PrP antibody 8H4 was used to RIP a subset of miRNAs in C2C12 myoblasts stably overexpressing WT PrP^C^ and C2C12 myoblasts (WT PrP^C^ RIP and Control RIP) when incubated with the differentiation medium for 4 days. Heat map shows the expression levels of miRNAs (left) and fold-enrichment compared to Inputs in C2C12 myoblasts (Control Input) and C2C12 myoblasts stably expressing WT PrP^C^ (WT PrP^C^ Input) (right). A subset of PrP^C^-bound miRNAs (red) is significantly enriched in WT PrP^C^ RIP upon differentiation for 4 days, compared to Control RIP and the two Inputs. **b**, Anti-PrP antibody-binding beads were used to IP endogenous PrP^C^ in C2C12 myoblasts differentiated for 4 days and then detected by Western blot with the anti-PrP antibody 8H4 and anti-GAPDH antibody, respectively. All blots also show the position of the molecular-weight markers. **c**, Volcano plot of 31 PrP^C^-bound miRNAs identified by small RNA-seq in differentiating C2C12 cells (control). Representative PrP^C^ RIP-enriched miRNAs, such as miR-486a-5p and miR-181b-5p, are highlighted in orange. **d**, Volcano plot of 51 PrP^C^-bound miRNAs identified by small RNA-seq in differentiating C2C12 cells stably expressing WT PrP^C^. Representative PrP^C^ RIP-enriched miRNAs, such as miR-214-3p and miR-204-5p, are highlighted in orange. Gray, PrP^C^ RIP-depleted miRNAs (**c**,**d**). **e**, Venn diagram of miRNAs enriched by PrP^C^ RIP in C2C12 cells stably expressing WT PrP^C^ versus miRNAs enriched by PrP^C^ RIP in C2C12 myoblasts (control). Group I (*n* = 21, Control RIP only), *n* = 10 (overlap), and group II (*n* = 41, WT PrP RIP only). **f**, RT−qPCR validation of depleted versus enriched miRNAs in differentiating C2C12 myoblasts (control) normalized to total input. **g**, RT−qPCR validation of depleted versus enriched miRNAs in differentiating C2C12 cells stably expressing WT PrP^C^ normalized to total input. **f**,**g**, U6 snRNA served as a negative control. Data are presented as means ± S.D. (*n* = 3 biologically independent measurements). **h**,**i**, The top six biological processes (BP) of Gene Ontology (GO) enrichment in the control cells (**h**) and the top eight BPs of GO enrichment in C2C12 cells stably overexpressing WT PrP^C^ (**i**). These significantly enriched pathways are related to cell differentiation and autophagy and may reflect the regulatory roles of miRNAs enriched by PrP^C^. **j**, Pulldown of endogenous PrP^C^ and excess PrP^C^ with WT and mutant biotin-labeled miR-214-3p or biotin-labeled miR-204-5p. Nonspecific miR-183-5p and miR-214-3p mutant were served as negative controls in biotinylated miRNA pulldown. Two GCAG sequences (orange) present in miR-214-3p, the predicted binding sites for PrP^C^, are mutated into two AAAA sequences (orange) in miR-214-3p mutant. PrP^C^ specifically interacts with miR-214-3p and miR-204-5p but does not interact with miR-183-5p and miR-214-3p mutant in both C2C12 cells stably expressing WT PrP^C^ and C2C12 cells during differentiation.

The secondary structure of miR-214-3p, an example of 41 miRNAs bound by PrP^C^, was predicted by RNAComposer^67^ (Extended Data Fig. 1a), and the structure of mouse PrP^C^ (PDB 1XYX)^68^ shows three α-helices (α1, α2, and α3) in its C-terminal domain (Extended Data Fig. 1b). We next used HDOCK, a protein−protein/nucleic acid protein docking web server by combining template-based and free docking^69^, to predict the binding sites of miR-214-3p for PrP^C^, and the molecular docking of the protein with miR-214-3p was performed using HDOCK (Extended Data Fig. 1b). The interface between PrP^C^ and miR-214-3p features two π-bonds between Tyr225 in PrP^C^ and GMP8 in miR-214-3p and a salt bridge between Arg228 in PrP^C^ and CMP9 in miR-214-3p (Extended Data Fig. 1c,d). Asp166, Gln167, Tyr168, Ser169, Asn170, Gln171, Val214, Thr 215, Tyr217, Gln218, Gln222, Tyr224, and Tyr225 in PrP^C^ form abundant hydrogen bonds with two GCAG sequences present in miR-214-3p (Extended Data Fig. 1d), which contribute to maintenance of the structure of the PrP^C^-miR-214-3p complex. Therefore, two GCAG sequences present in miR-214-3p are the predicted binding sites for PrP^C^ and were mutated into two AAAA sequences in miR-214-3p mutant. We then pulled down endogenous PrP^C^ and excess PrP^C^ with WT and mutant biotin-labeled miR-214-3p or biotin-labeled miR-204-5p from C2C12 cell extracts, using nonspecific miR-183-5p and miR-214-3p mutant as negative controls (Fig. 4j). Of note, miR-214-3p and miR-204-5p were able to capture both endogenous PrP^C^ and excess PrP^C^, and mutating the GCAG sequences in miR-214-3p completely abolished its ability to capture PrP^C^; endogenous PrP^C^ specifically interacted with miR-214-3p and miR-204-5p but did not interact with miR-183-5p and miR-214-3p mutant in control C2C12 cells during differentiation; excess PrP^C^ also specifically interacted with miR-214-3p and miR-204-5p but did not interact with miR-183-5p and miR-214-3p mutant in C2C12 cells stably expressing WT PrP^C^ during differentiation (Fig. 4j). Together, these results demonstrate PrP^C^ selectively binds to a subset of miRNAs such as miR-214-3p and miR-204-5p during myoblast differentiation.

### MiR-214-3p inhibits ATG5-dependent autophagy and myoblast differentiation via interaction with PrP^C^

Given that PrP^C^ selectively binds to miR-214-3p and miR-204-5p during myoblast differentiation (Fig. 4), we predicted that these two miRNAs might regulate autophagy and differentiation of skeletal muscle cells via interaction with PrP^C^. We next used miRNA reporter gene assays and Western blotting to test this hypothesis. Using TargetScan and miRanda algorithms^70^, we focused on two candidate target genes, *ATG5* for miR-214-3p and *LC3B* for miR-204-5p, and tested their functional responses. We used miR-214-3p and miR-204-5p reporter gene assays, in which miR-214-3p represses the wild-type 3’ end of the untranslated region (WT 3’UTR) of the gene *ATG5* associated with luciferase (Fig. 5a) and miR-204-5p represses the WT 3’UTR of the gene *LC3B* associated with luciferase (Fig. 5b). The mutant versions of the 3’UTR (Mut 3’UTR), in which the miR-214-3p binding site (Fig. 5a) and the miR-204-5p binding site (Fig. 5b) were mutated, were served as negative controls in miRNA reporter gene assays. Compared with transfection of control (NC) in C2C12 myoblasts KO for PrP^C^, transfection of miR-214-3p mimic (Fig. 5a) or miR-204-5p mimic (Fig. 5b) at 10 μM significantly inhibited the relative luciferase activity of the reporter (*p* = 0.001 or 0.035) but did not significantly change that of the mutant version of the reporter (*p* = 0.27 or 0.93) in this cell line. Importantly, compared with transfection of NC in C2C12 myoblasts stably expressing WT PrP^C^, transfection of miR-214-3p mimic (Fig. 5a) or miR-204-5p mimic (Fig. 5b) at 10 μM more significantly inhibited the relative luciferase activity of the reporter (*p* = 0.00027 or 0.001) but did not significantly change that of the mutated reporter (*p* = 0.47 or 0.37) in the C2C12 myoblasts. Notably, compared with that in C2C12 myoblasts KO for PrP^C^, excess PrP^C^ significantly enhanced not only the binding affinity of miR-214-3p towards its downstream target, the autophagy marker ATG5 (Fig. 5a) (*p* = 0.00021), but also the binding affinity of miR-204-5p towards its downstream target, the autophagy marker LC3B (Fig. 5b) (*p* = 0.0047), in C2C12 myoblasts stably expressing WT PrP^C^ when incubated with the differentiation medium for 4 days. To gain a quantitative understanding of how miR-214-3p regulates skeletal muscle cell autophagy, we performed Western blot analysis for ATG5 and the myogenic differentiation marker MyHC after transfection with NC or miR-214-3p inhibitor at 10 μM in the above cell lines (Fig. 5c). Upon differentiation for 4 days, the relative amounts of ATG5 (Fig. 5d) and MyHC (Fig. 5e) in the cell lysates from C2C12 myoblasts stably expressing WT PrP^C^ or KO for PrP^C^ transfected with 10 μM miR-214-3p inhibitor were significantly higher than those in the cell lysates from the above cell lines transfected with NC. Thus, compared with transfection of NC in C2C12 myoblasts KO for PrP^C^ and C2C12 myoblasts stably expressing WT PrP^C^, transfection of 10 μM miR-214-3p inhibitor significantly promoted ATG5-dependent autophagy and myoblast differentiation in the above cell lines (Fig. 5c−e), suggesting that miR-214-3p significantly inhibits both ATG5-dependent autophagy and myoblast differentiation via specific interaction with PrP^C^. Importantly, after transfection with miR-214-3p inhibitor, excess PrP^C^ significantly inhibited both ATG5-dependent autophagy and myoblast differentiation in differentiating C2C12 cells stably expressing WT PrP^C^, compared to those in differentiating C2C12 myoblasts KO for PrP^C^ (Fig. 5d,e). To gain a quantitative understanding of how miR-204-5p regulates skeletal muscle cell autophagy, we performed Western blot analysis for LC3B and MyHC after transfection with NC or miR-204-5p inhibitor at 10 μM in the above cell lines (Fig. 5f). Upon differentiation for 4 days, transfection of 10 μM miR-204-5p inhibitor significantly promoted LC3B-dependent autophagy in C2C12 myoblasts KO for PrP^C^ and C2C12 myoblasts stably expressing WT PrP^C^, compared to transfection of NC in the above cell lines (Fig. 5g). Importantly, PrP^C^ deficiency completely blocked muscle cell differentiation after transfection with NC or miR-204-5p inhibitor at 10 μM in C2C12 myoblasts KO for PrP^C^ upon differentiation for 4 days (Fig. 5h). Compared with transfection of NC in C2C12 myoblasts stably expressing WT PrP^C^, however, transfection of miR-204-5p inhibitor significantly promoted myoblast differentiation in the above cell line (Fig. 5h), suggesting that miR-204-5p, another PrP^C^-bound miRNA, significantly inhibits myoblast differentiation. These results demonstrate that miR-214-3p significantly inhibits ATG5-dependent autophagy and myoblast differentiation via specific interaction with PrP^C^. Similarly, miR-204-5p significantly inhibits LC3B-dependent autophagy and myoblast differentiation via specific interaction with PrP^C^.

**Fig. 5.**
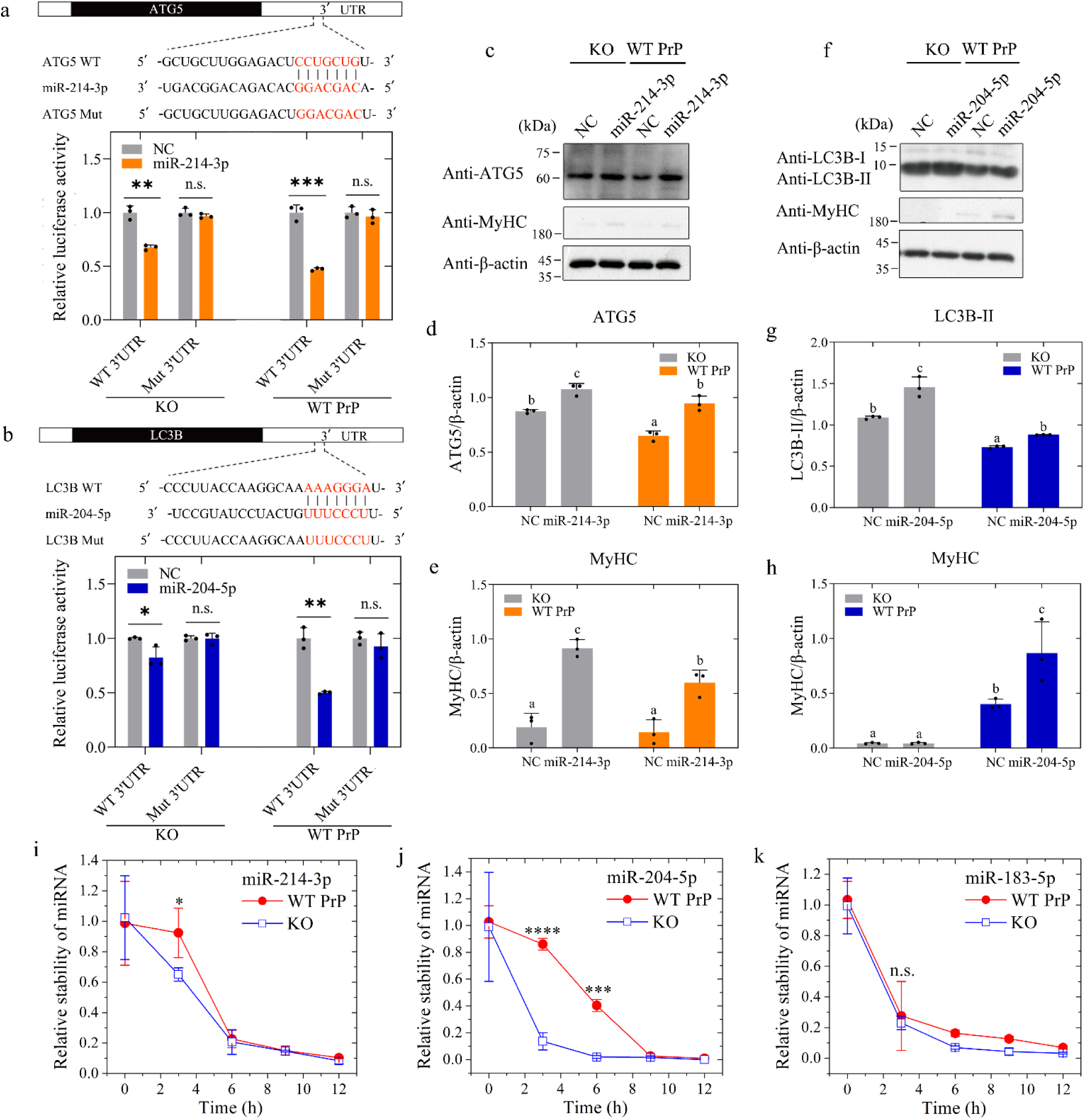
MiR-214-3p significantly inhibits both ATG5-dependent autophagy and myoblast differentiation via specific interaction with PrP^C^. **a**,**b**, Compared with that in C2C12 myoblasts KO for PrP^C^, excess PrP^C^ significantly enhances not only the binding affinity of miR-214-3p towards its downstream target, the autophagy marker ATG5 (**a**) (*p* = 0.00021), but also the binding affinity of miR-204-5p towards its downstream target, the autophagy marker LC3B (**b**) (*p* = 0.0047), in C2C12 myoblasts stably overexpressing WT PrP^C^ when incubated with the differentiation medium for 4 days. We used miR-214-3p and miR-204-5p reporter gene assays, in which miR-214-3p represses the wild-type 3’ end of untranslated region (WT 3’UTR) of the gene *ATG5* associated with luciferase (**a**) and miR-204-5p represses the WT 3’UTR of the gene *LC3B* associated with luciferase (**b**). The mutant versions of the 3’UTR (Mut 3’UTR), in which the miR-214-3p binding site (**a**) and the miR-204-5p binding site (**b**) were mutated, were served as negative controls in miRNA reporter gene assays. Each pair of reporters was assayed in response to specific miRNA mimics in the above cell lines. Compared with transfection of control (NC) (gray) in C2C12 myoblasts KO for PrP^C^, transfection of miR-214-3p mimic (orange) (**a**) or miR-204-5p mimic (blue) (**b**) at 10 μM significantly inhibits the relative luciferase activity of the reporter (*p* = 0.001 or 0.035) but does not significantly change that of the mutant version of the reporter (*p* = 0.27 or 0.93) in this cell line. Importantly, compared with transfection of NC in C2C12 myoblasts stably expressing WT PrP^C^, transfection of miR-214-3p mimic (orange) (**a**) or miR-204-5p mimic (blue) (**b**) at 10 μM more significantly inhibits the relative luciferase activity of the reporter (*p* = 0.00027 or 0.001) but does not significantly change that of the mutated reporter (*p* = 0.47 or 0.37) in the C2C12 myoblasts. **c**, Western blot for ATG5 and the myogenic differentiation marker MyHC after transfection with NC or miR-214-3p inhibitor at 10 μM in the above cell lines upon differentiation for 4 days. The cell lysates from the above cells were probed by the anti-ATG5 antibody, anti-MyHC antibody, and anti-β-actin antibody, respectively. **d**,**e**, The relative amount of ATG5 (**d**) or MyHC (**e**) in the above cell lines (solid black circles shown in scatter plots) was determined as a ratio of the density of ATG5 or MyHC bands over the density of β-actin band in cell lysates and expressed as the mean ± S.D. (with error bars) of values obtained in three independent experiments. Compared with transfection of NC in C2C12 myoblasts KO for PrP^C^ (gray) and C2C12 myoblasts stably expressing WT PrP^C^ (orange), transfection of 10 μM miR-214-3p inhibitor significantly promotes ATG5-dependent autophagy and myoblast differentiation in the above cell lines. **f**, Western blot for LC3B and MyHC after transfection with NC or miR-204-5p inhibitor at 10 μM in the above cell lines upon differentiation for 4 days. The cell lysates from the above cells were probed by the anti-LC3B antibody, anti-MyHC antibody, and anti-β-actin antibody, respectively. All blots also show the position of the molecular-weight markers. **g**,**h**, The relative amount of LC3B-II (**g**) or MyHC (**h**) in the above cell lines (solid black circles shown in scatter plots) was determined as a ratio of the density of LC3B-II or MyHC bands over the density of β-actin band in cell lysates and expressed as the mean ± S.D. (with error bars) of values obtained in three independent experiments. **g**, Transfection of 10 μM miR-204-5p inhibitor significantly promotes LC3B-dependent autophagy in C2C12 myoblasts KO for PrP^C^ (gray) and C2C12 myoblasts stably expressing WT PrP^C^ (blue), compared to transfection of NC in the above cell lines. **h**, PrP^C^ deficiency completely blocks muscle cell differentiation after transfection with NC or miR-204-5p inhibitor at 10 μM in C2C12 myoblasts KO for PrP^C^ (gray) upon differentiation for 4 days. Compared with transfection of NC in C2C12 myoblasts stably expressing WT PrP^C^ (blue), however, transfection of miR-204-5p inhibitor significantly promotes myoblast differentiation in the above cell line. **i**–**k**, Compared with that in C2C12 myoblasts KO for PrP^C^ (blue), excess PrP^C^ significantly enhances the relative stability of a subset of miRNAs, miR-214-3p (*p* = 0.048 at incubation time of 3 h) (**i**) and miR-204-5p (*p* = 0.000086 and 0.000136 at incubation time of 3 and 6 h, respectively) (**j**), in C2C12 myoblasts stably expressing WT PrP^C^ (red) during myoblast differentiation, but does not significantly change the relative stability of nonspecific miR-183-5p (*p* = 0.74 and 0.21 at 3 and 6 h, respectively) (**k**), served as a negative control, in this cell line. C2C12 myoblasts stably expressing WT PrP^C^ and C2C12 myoblasts KO for PrP^C^ were incubated with the differentiation medium for 4 days and then treated with 20 mg/ml α-amanitin, an inhibitor of RNA polymerase II. The above cell lines were lysed every 3 h for up to 12 h. The relative stability of miRNAs in the above cell lines was determined as the relative enrichment (to U6 snRNA) of miRNAs by RT−qPCR and expressed as the mean ± S.D. (with error bars) of values obtained in three independent experiments. The relative stability of miRNAs is decreased with the increase of incubation time of α-amanitin. **a**,**b**,**i**–**k**, Statistical analyses were performed using a two-tailed unpaired *t* test. Values of *p* < 0.05 indicate statistically significant differences. The following notation is used throughout: \**p* < 0.05; \*\**p* < 0.01; \*\*\**p* < 0.001; and \*\*\*\**p* < 0.0001 relative to controls. n.s., no significance. **d**,**e**,**g**,**h**, One-way ANOVA and multiple comparisons were performed by SPSS 19.0 and different letters indicate significant differences at the level of *p* < 0.05.

### Overexpressed PrP^C^ is colocalized with miR-214-3p in the skeletal muscle of myopathy patients

PrP^C^ dysfunction and miRNA overexpression are both observed in the skeletal muscle of myopathy patients^19,22,37^. Interestingly, we found that miR-214-3p, an example of 41 miRNAs bound by PrP^C^, significantly inhibited autophagy and differentiation of skeletal muscle cells (Figs. 4 and 5a−h). Based on these observations, we hypothesized that PrP^C^ is colocalized with miR-214-3p in the skeletal muscle of myopathy patients. To test this hypothesis, we took confocal images of frozen skeletal muscle sections from six myopathy patients and four age-matched controls (Fig. 1a,b). The frozen skeletal muscle sections, in which miR-214-3p was detected by FISH (green) using an FAM-labeled miR-214-3p probe, were immunostained with the anti-PrP antibody 8H4 (red), stained with DAPI (blue), and visualized by confocal microscopy. Excess PrP^C^ (red) mainly located in the cytoplasm, abundant miR-214-3p (green dots), and the co-localization of excess PrP^C^ with miR-214-3p (yellow dots in the merged images) were clearly observed in the skeletal muscle of these myopathy patients (Fig. 1a). In sharp contrast, PrP^C^ overexpression, miR-214-3p expression, and the co-localization of PrP^C^ with miR-214-3p were all not observed in the skeletal muscle of those controls (Fig. 1b). Thus, overexpressed PrP^C^ was colocalized with miR-214-3p in the skeletal muscle of six myopathy patients (Fig. 1a). In sharp contrast, such a phenomenon was not observed in the age-matched controls (Fig. 1b). To further test this hypothesis, we next used C2C12 mouse myoblasts (control) and C2C12 myoblasts stably overexpressing WT PrP^C^ upon differentiation for 4 days, in which miR-214-3p was detected by FISH (green) using an FAM-labeled miR-214-3p probe. The above two cell lines were immunostained with the anti-PrP antibody 8H4 (red), stained with DAPI (blue), and visualized by confocal microscopy (Fig. 1e). Excess PrP^C^ (red) mainly located in the cytoplasm, abundant miR-214-3p (green dots), and the co-localization of excess PrP^C^ with miR-214-3p (yellow dots in the merged images) were clearly observed in differentiating C2C12 cells stably expressing WT PrP^C^ (Fig. 1e). Both miR-214-3p expression and the co-localization of endogenous PrP^C^ with miR-214-3p (the merged images), however, were not observed in differentiating C2C12 cells (control) (Fig. 1e). Thus, overexpressed PrP^C^ was colocalized with miR-214-3p in differentiating C2C12 cells stably expressing WT PrP^C^ (Fig. 1e). In sharp contrast, such a phenomenon was not observed in control C2C12 cells (Fig. 1e). Moreover, we wanted to know whether PrP^C^ could modulate the stability of mature miR-214-3p in C2C12 myoblasts (Fig. 5i−k). C2C12 myoblasts stably expressing WT PrP^C^ or KO for PrP^C^ were incubated with the differentiation medium for 4 days and then treated with 20 mg/ml α-amanitin, an inhibitor of RNA polymerase II. Compared with that in C2C12 myoblasts KO for PrP^C^, excess PrP^C^ significantly enhanced the relative stability of miR-214-3p and miR-204-5p (Fig. 5i,j), two examples of 41 miRNAs bound by PrP^C^, in C2C12 myoblasts stably expressing WT PrP^C^ during myoblast differentiation, but did not significantly change the relative stability of nonspecific miR-183-5p (Fig. 5k), a negative control, in this cell line. Together, the data showed that under pathological conditions, overexpressed PrP^C^ is colocalized with miR-214-3p in skeletal muscle cells to increase the stability of mature miR-214-3p.

### PrP^C^ selectively recruits a subset of miRNAs into phase-separated condensates, which in turn greatly enhances the LLPS of PrP^C^

Given that PrP^C^ specifically interacts with a subset of miRNAs such as miR-214-3p and miR-204-5p during myoblast differentiation (Fig. 4), we predicted that miR-214-3p and miR-204-5p might regulate LLPS of PrP^C^ via interaction with the protein. We next used confocal microscopy and fluorescence recovery after photobleaching (FRAP)^48,49,51,52,54−56^ to test this hypothesis. Bacterial purified WT mouse PrP^C^, labeled by 5(6)-carboxy-tetramethylrhodamine *N*-succinimidyl ester (TAMRA, red fluorescence) and incubated with 1 × PBS (pH 7.4) on ice, underwent LLPS in vitro and formed protein condensates (Extended Data Fig. 2). PrP^C^ formed abundant liquid droplets and protein condensates formed by PrP^C^ became much larger in the presence of miR-214-3p or miR-204-5p of low concentrations, compared to those in the absence of miRNA (Fig. 6a). Low concentrations of miR-214-3p and miR-204-5p dramatically promoted in vitro LLPS of PrP^C^ (Fig. 6a). In sharp contrast, low concentrations of nonspecific miR-183-5p, a negative control, only mildly enhanced in vitro LLPS of PrP^C^ (Fig. 6a). To further test this hypothesis, we next took fluorescence images of in vitro phase-separated PrP^C^ droplets with three miRNAs (Fig. 6b). 50 μM WT PrP^C^, labeled by TAMRA (red fluorescence) and incubated with 1 × PBS (pH 7.4) containing 10 μM FAM-labeled miRNA (green fluorescence) on ice, also underwent LLPS in vitro (Fig. 6b). PrP^C^ de-mixed droplets (red; Merge: yellow) fused with droplets of miR-214-3p or those of miR-204-5p (green) were clearly observed by confocal microscopy, with excitation at 561 nm and 488 nm, respectively (Fig. 6b). Importantly, PrP^C^ selectively recruited and concentrated a subset of miRNAs into phase-separated condensates (Fig. 6b). The recruitment ability of PrP^C^ on miR-214-3p and miR-204-5p (green; Merge: yellow) was much stronger than that on two PrP^C^-unbound miRNAs (negative controls), miR-183-5p and mutant miR-214-3p (light green; Merge: orange) (Fig. 6b). Addition of miR-214-3p or miR-204-5p of low concentrations dramatically promoted but the addition of miR-183-5p or miR-214-3p mutant did not dramatically enhance the phase separation (Fig. 6a) and droplet fusion ability (Fig. 6b) of PrP^C^. Together, the data showed that a subset of miRNAs recruited by PrP^C^ forms a positive feed-forward loop with PrP^C^ to enhance in vitro LLPS of PrP^C^ greatly.

**Fig. 6.**
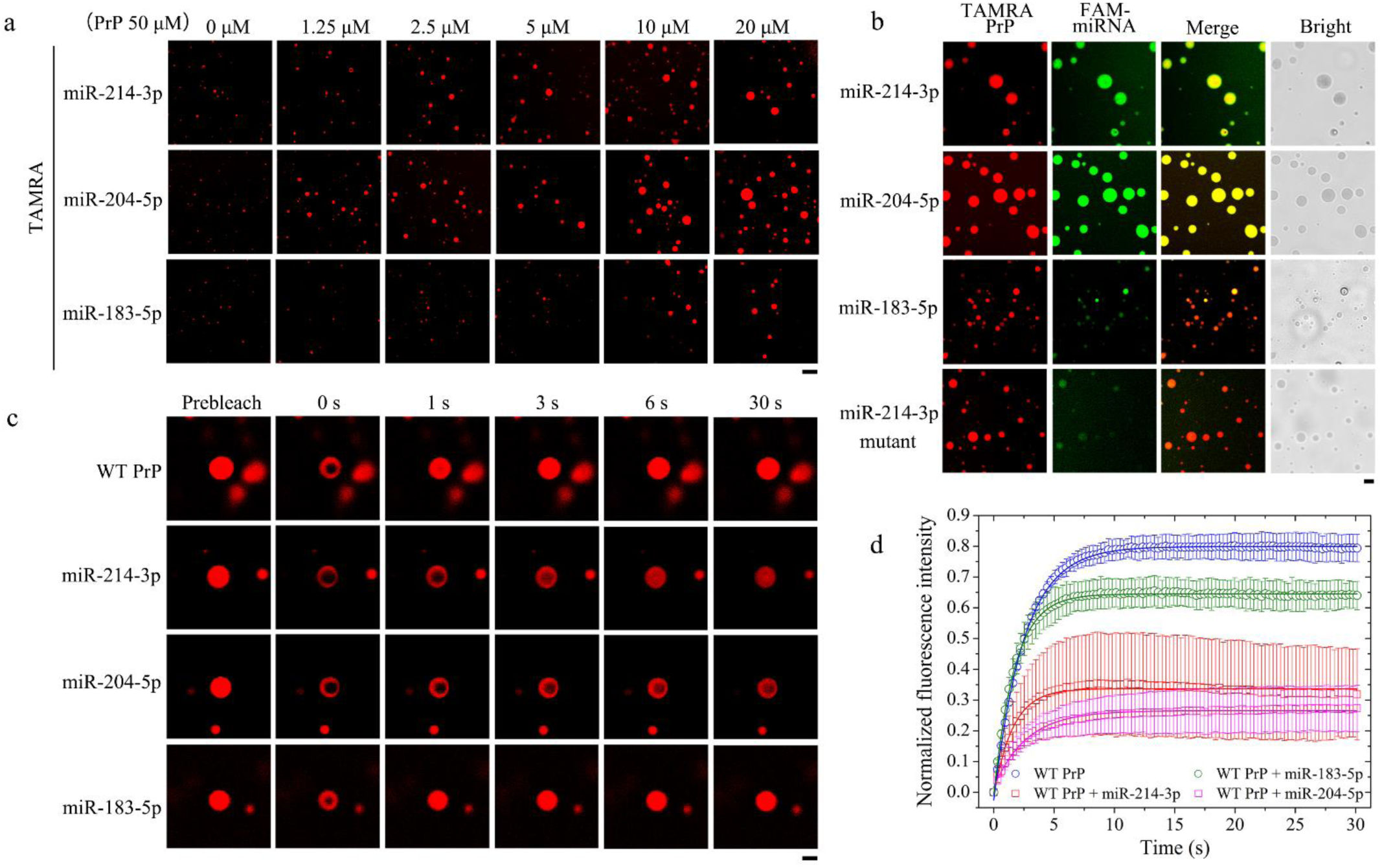
PrP^C^ selectively recruits a subset of miRNAs into phase-separated condensates, which in turn greatly enhances in vitro LLPS of PrP^C^. **a**, Regulation of PrP^C^ LLPS by three miRNAs. Samples (50 μM) of bacterial purified wild-type mouse PrP^C^ (WT PrP^C^) were labeled by TAMRA (red fluorescence) and incubated with 1 × PBS (pH 7.4) containing 0, 1.25, 2.5, 5, 10 or 20 μM miRNA on ice to induce LLPS for 5 min. Liquid droplets of PrP^C^ (protein condensates) were observed by confocal microscopy, with excitation at 561 nm. Low concentrations of miR-214-3p and miR-204-5p dramatically promote in vitro LLPS of PrP^C^. In sharp contrast, low concentrations of nonspecific miR-183-5p, served as a negative control in the in vitro phase separation assay, only mildly enhances in vitro LLPS of PrP^C^. Scale bar, 7.5 nm. **b**, Fluorescence images of in vitro phase-separated PrP^C^ droplets with three miRNAs. Samples (50 μM) of WT PrP^C^ were labeled by TAMRA (red fluorescence) and incubated with 1 × PBS containing 10 μM FAM-labeled miRNA (green fluorescence) on ice to induce LLPS for 5 min. PrP^C^ de-mixed droplets (red; Merge: yellow) fused with droplets of miR-214-3p or those of miR-204-5p (green) were clearly observed by confocal microscopy, with excitation at 561 nm and 488 nm, respectively. The right column shows the brightfield images for PrP^C^ LLPS in the presence of miRNA. Importantly, PrP^C^ selectively recruits and concentrates a subset of miRNAs into phase-separated condensates. The recruitment ability of PrP^C^ on miR-214-3p and miR-204-5p (green; Merge: yellow) is much stronger than that on two PrP^C^-unbound miRNAs, miR-183-5p and mutant miR-214-3p (light green; Merge: orange). Mutant miR-214-3p, in which two GCAG sequences predicted as binding sites for PrP^C^ were mutated into two AAAA sequences, was served as another negative control in the in vitro phase separation assay. Addition of miR-214-3p or miR-204-5p of low concentrations dramatically promotes but the addition of miR-183-5p or miR-214-3p mutant does not dramatically enhance the phase separation (**a**) and droplet fusion ability (**b**) of PrP^C^. Scale bar, 2.5 nm. **c**,**d**, miR-214-3p and miR-204-5p of low concentrations decrease fluorescence recovery and modulate the liquid nature of PrP^C^ droplets in vitro. **c**, Representative examples of in vitro FRAP analysis of phase-separated PrP^C^ droplets with/without miRNA. FRAP analysis on the selected liquid droplets of 50 μM WT PrP^C^ labeled by TAMRA (red fluorescence) before (prebleach), during (0 s), and after photobleaching (1, 3, 6 and 30 s, respectively). The internal photobleaching is marked by a black square. These samples were incubated with 1 × PBS containing 10 μM miRNA on ice to induce LLPS for 10 min. Scale bar, 2.5 nm. **d**, Normalized kinetics of fluorescence recovery data of WT PrP^C^ (open blue circle), WT PrP^C^ + miR-214-3p (open red square), WT PrP^C^ + miR-204-5p (open magenta square), and WT PrP^C^ + miR-183-5p (open olive circle) obtained from FRAP intensity. The normalized fluorescence intensity is expressed as the mean ± S.D. (with error bars) of values obtained in three independent experiments. The solid lines show the best single exponential fit for the fluorescence intensity-time curves. FRAP of in vitro phase-separated PrP^C^ droplets without miRNA or with a negative control miR-183-5p revealed a (82.5 ± 0.3)% or (65.1 ± 0.3)% fluorescence recovery within 30 s. In sharp contrast, FRAP of in vitro phase-separated PrP^C^ droplets coacervated with miR-214-3p or miR-204-5p revealed a much lower fluorescence recovery, (34.5 ± 0.8)% or (24.9 ± 0.3)%, within 30 s. All FRAP experiments were repeated three times and the results were reproducible.

We then investigated and evaluated the dynamics of in vitro phase-separated droplets of PrP^C^ with/without miRNA by FRAP (Fig. 6c,d). FRAP of phase-separated PrP^C^ droplets without miRNA or with a negative control miR-183-5p revealed a (82.5 ± 0.3)% or (65.1 ± 0.3)% fluorescence recovery within 30 s (Fig. 6d). In sharp contrast, FRAP of phase-separated PrP^C^ droplets coacervated with miR-214-3p or miR-204-5p revealed a much lower fluorescence recovery, (34.5 ± 0.8)% or (24.9 ± 0.3)%, within 30 s (Fig. 6d). According to Fig. 6c,d, miR-214-3p and miR-204-5p reduced the fluorescence recovery. It means miR-214-3p and miR-204-5p decrease the fluidity of LLPS condensates, possibly because these miRNAs could modulate liquid-to-solid transitions in phase-separated PrP^C^ condensates. The above experiments help drive the narrative that miR-214-3p and miR-204-5p of low concentrations decrease fluorescence recovery and modulate the liquid nature of PrP^C^ droplets in vitro.

Altogether, these data strongly suggest that the interactions between PrP^C^ and miR-214-3p or miR-204-5p control liquidity, and miR-214-3p and miR-204-5p reduce PrP^C^ mobility via specific interaction with PrP^C^. Therefore, miR-214-3p and miR-204-5p are key factors in modulating PrP^C^ liquid-phase condensation.

### PrP^C^ recruits miR-214-3p into phase-separated condensates in living skeletal muscle cells, which in turn promotes pathological aggregation of PrP

Given that PrP^C^ selectively recruits a subset of miRNAs into phase-separated condensates, which in turn greatly enhances in vitro LLPS of PrP^C^ (Fig. 6), we predicted that miR-214-3p, an example of these miRNAs, might regulate in vivo LLPS of PrP^C^ and the subsequent PrP aggregation in skeletal muscle cells. PrP^C^ contains an IDR in its N-terminal domain^7,48−52,54−56,68^. We next employed an optogenetic tool that uses a blue light (488-nm laser) to activate IDR-mediated LLPS of proteins in living cells^71−73^ and immunogold electron microscopy^74−76^ to test this hypothesis. We used this ‘‘optoDroplet’’ system (Fig. 7a) to take time-lapse images of living C2C12 cells expressing WT PrP^C^-optoIDR construct, which contains PrP_1-37_ IDR (residues 1–37) linked to mCherry fluorescent protein and Cry2 and then linked to PrP_38-230_ IDR (residues 38–230), and living C2C12 cells transfected with optoIDR plasmids (IDR-mCherry-Cry2) (Fig. 7b,c). We also took time-lapse images (0–100 s) of a C2C12 cell expressing the WT PrP^C^-optoIDR construct (Fig. 7d,e). Notably, PrP^C^ underwent light-activated LLPS and formed abundant liquid droplets (white puncta) in the cytoplasm of living C2C12 cells (Fig. 7c,d). Two small liquid condensates gradually fused into one larger liquid droplet in living C2C12 cells (Fig. 7e and Supplementary Movie 1). To test the first half of this hypothesis, we took live-cell images of PrP^C^-miRNA speck formation when 10 μM FAM-labeled miRNA (green) was transfected into C2C12 cells stably expressing mCherry (red) or WT PrP^C^-mCherry (red) (Fig. 7f). PrP^C^ was colocalized with miR-214-3p in phase-separated condensates (Fig. 7f and Supplementary Movies 2−5). PrP^C^ de-mixed droplets (red; Merge: yellow) fused with droplets of miR-214-3p (green), and the co-localization of PrP^C^ with miR-214-3p (yellow puncta in the merged images) in phase-separated condensates was clearly observed by confocal microscopy, with excitation at 561 nm and 488 nm, respectively (Fig. 7f). Together, the data showed that PrP^C^ recruited and concentrated miR-214-3p into phase-separated condensates in living skeletal muscle cells.

**Fig. 7.**
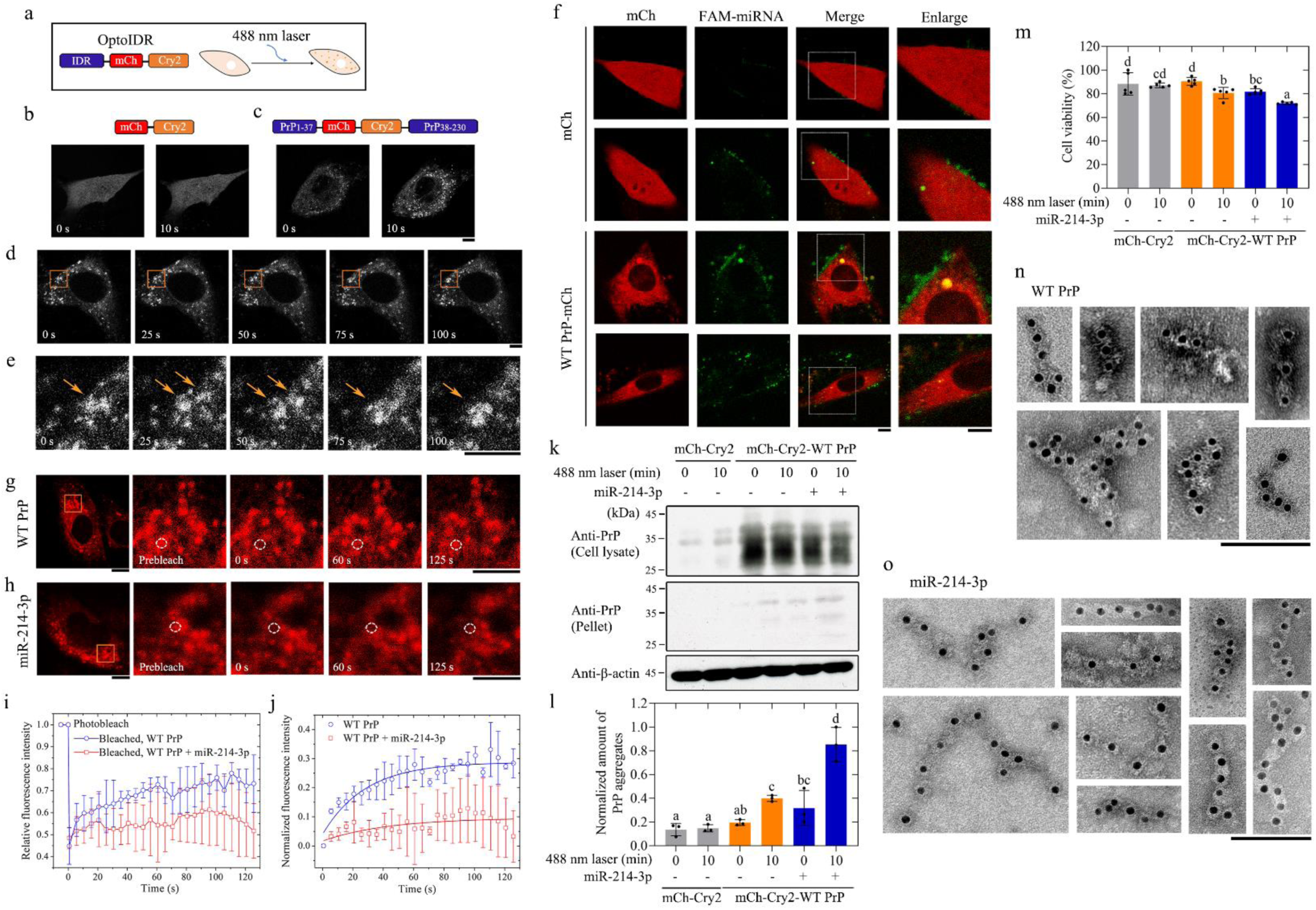
PrP^C^ recruits miR-214-3p into phase-separated condensates in living skeletal muscle cells, which in turn promotes pathological aggregation of PrP. **a**, Rapid light-dependent clustering of intrinsically disordered region (IDR) (blue) fused Cry2 (orange) and a schematic diagram of the optogenetic platform (optoDroplets) consisting of the N-terminal IDR fused to mCherry fluorescent protein (mCh) (red) and Cry2 (left). Blue light activation of optoIDRs using a 488-nm excitation laser (right). **b**,**c**, Time-lapse images of living C2C12 cells expressing WT PrP^C^-optoIDR construct containing PrP_1-37_ IDR (residues 1–37) (blue) linked to mCh (red) and Cry2 (orange) and then linked to PrP_38-230_ IDR (residues 38–230) (blue) (**c**). mCh-Cry2 fusion alone was used as a control (**b**). **d**,**e**, Time-lapse images (0–100 s) of a C2C12 cell expressing the WT PrP^C^-optoIDR construct. The enlarged regions (**e**) show 25-fold enlarged images from (**d**). PrP^C^ undergoes light-activated LLPS and forms abundant liquid droplets (white puncta) in the cytoplasm of living C2C12 cells (**c**,**d**), which was observed by confocal microscopy, with excitation at 561 nm. **e**, Two small liquid droplets gradually fuse into one larger liquid droplet in the enlarged region (highlighted by using orange arrows) of living C2C12 cells (see also Supplementary Movie 1). **b**–**e**, Cells were cultured for 24.5 h and subjected to laser excitation every 2 s for the indicated time. Scale bars, 7.5 μm. **f**, Representative live-cell images of PrP^C^-miRNA speck formation when 10 μM FAM-labeled miRNA (green fluorescence) was transfected into C2C12 cells stably expressing mCh (red fluorescence) (the first and second rows) or WT PrP^C^-mCh (red) (the third and fourth rows), among ≥ 10 cells, showing that PrP^C^ is colocalized with miR-214-3p in phase-separated condensates (see also Supplementary Movies 2–5). The enlarged regions (right) show 4-fold enlarged images from the merged images. PrP^C^ de-mixed droplets (red; Merge: yellow) fused with droplets of miR-214-3p (green), and the co-localization of PrP^C^ with miR-214-3p (yellow puncta in the merged images) in phase-separated condensates were clearly observed by confocal microscopy, with excitation at 561 nm and 488 nm, respectively. Scale bars, 7.5 μm. **g**–**j**, transfection of 10 μM miR-214-3p decreases fluorescence recovery and modulates the liquid nature of PrP^C^ droplets in vivo. **g**,**h**, Representative examples of FRAP analysis of light-activated phase-separated PrP^C^ droplets without or with miRNA (**g** or **h**) in the enlarged region of living C2C12 cells (see also Supplementary Movies 6 and 7). The enlarged regions show 25-fold enlarged images from the left. In vivo FRAP analysis on the selected liquid droplets of WT PrP^C^ (red fluorescence) before (prebleach), during (0 s), and after photobleaching (60 and 125 s, respectively). The dashed white circle highlights the punctum undergoing targeted bleaching. Cells were cultured for 24 h, transfected without or with 10 μM miR-214-3p for 30 min in a growing medium, and subjected to laser excitation for 10 s. Images were then captured by confocal microscopy, with excitation at 561 nm. Scale bars, 7.5 μm. **b**–**e**,**g**,**h**, All cells here are at similar expression levels and activated under identical conditions. **i**, Quantification of FRAP data for WT PrP^C^ puncta without or with miRNA (*n* = 3), as in **g**,**h**. Time 0 indicates the start of recovery after photobleaching. **j**, Normalized kinetics of fluorescence recovery data of WT PrP^C^ puncta (open blue circle) and WT PrP^C^ puncta + miR-214-3p (open red square) in living C2C12 cells. The normalized fluorescence intensity is expressed as the mean ± S.D. (with error bars) of values obtained in three independent experiments. The solid lines show the best single exponential fit for the fluorescence intensity-time curves. FRAP of in vivo phase-separated PrP^C^ droplets without miRNA revealed a (24.8 ± 1.8)% fluorescence recovery within 125 s. In sharp contrast, FRAP of in vivo phase-separated PrP^C^ droplets coacervated with 10 μM miR-214-3p revealed a much lower fluorescence recovery, (8.0 ± 1.9)%, within 125 s. All FRAP experiments were repeated three times and the results were reproducible. **k**–**m**, Transfection of 10 μM miR-214-3p significantly promotes light-activated pathological aggregation of PrP and significantly increase PrP toxicity in C2C12 myoblasts stably expressing WT PrP^C^. C2C12 cells stably overexpressing mCh-Cry2-WT PrP^C^ (the right four lanes in **k** and the right four bars in **l** or **m**) were cultured until confluence was reached 85% (**k**,**l**) or 80% (**m**), then transfected without (-) or with (+) 10 μM miR-214-3p for 30 min in a growing medium, and cultured for 12 h after activation by 488-nm laser for 10 min, using C2C12 myoblasts stably overexpressing mCh-Cry2 (the left two lanes in **k** and the left two bars in **l** or **m**) as a control and cells not activated by 488-nm laser as another control. **k**, Then the sarkosyl-insoluble pellets from the above cells were probed by the anti-PrP antibody 8H4, and the corresponding cell lysates were probed using 8H4 and anti-β-actin antibody, respectively. All blots also show the position of the molecular-weight markers. **l**, The normalized amount of insoluble PrP aggregates in the above cell lines (solid black circles shown in scatter plots) was determined as a ratio of the density of insoluble PrP aggregate bands over that of the total PrP bands in cell lysates and expressed as the mean ± S.D. (with error bars) of values obtained in three independent experiments. **m**, Then the above cells were detected by MTT reduction assay. The cell viability (%) (solid black circles shown in scatter plots) are expressed as the mean ± S.D. (with error bars) of values obtained in five independent experiments. **l**,**m**, One-way ANOVA and multiple comparisons were performed by SPSS 19.0 and different letters indicate significant differences at the level of *p* < 0.05. **n**,**o**, Immunogold electron microscopy of PrP fibrils purified from C2C12 myoblasts stably expressing WT PrP^C^. The cells were cultured until confluence was reached 85%, then transfected without (**n**) or with (**o**) 10 μM miR-214-3p for 30 min in a growing medium, cultured for 12 h after activation by 488-nm laser for 10 min, and labeled by gold particles conjugated with the anti-PrP antibody 8H4. The skeletal muscle cells transfected with 10 μM miR-214-3p produced much more amyloid fibrils than those transfected without miRNA, which were clearly observed by immunogold electron microscopy. Scale bars, 100 nm.

We then investigated and evaluated the dynamics of in vivo phase-separated droplets of PrP^C^ with/without miRNA by FRAP (Fig. 7g−j). PrP^C^ and miR-214-3p puncta exhibit features characteristic of liquid-like condensates (Fig. 7g,h and Supplementary Movies 6 and 7). FRAP of phase-separated PrP^C^ droplets without miRNA revealed a (24.8 ± 1.8)% fluorescence recovery within 125 s (Fig. 7j). In sharp contrast, FRAP of phase-separated PrP^C^ droplets coacervated with 10 μM miR-214-3p revealed a much lower fluorescence recovery, (8.0 ± 1.9)%, within 125 s (Fig. 7j). The above experiments help drive the narrative that transfection of 10 μM miR-214-3p decreases fluorescence recovery and modulates the liquid nature of PrP^C^ droplets in vivo. These data once again suggest that the interaction between PrP^C^ and miR-214-3p controls liquidity, and miR-214-3p reduces PrP^C^ mobility via specific interaction with PrP^C^.

To test the second half of this hypothesis, we used C2C12 cells stably expressing mCherry-Cry2-WT PrP^C^, which were cultured until confluence was reached 85% (Fig. 7k,l) or 80% (Fig. 7m), transfected without or with 10 μM miR-214-3p for 30 min, and cultured for 12 h after activation by 488-nm laser for 10 min, using C2C12 myoblasts stably expressing mCherry-Cry2 as a control. The sarkosyl-insoluble pellets from the above cells were probed by the anti-PrP antibody 8H4, and the corresponding cell lysates were probed using 8H4 and anti-β-actin antibody, respectively (Fig. 7k). The above cells were also detected by 3-(4,5-dimethylthiazol-2-yl)-2,5-diphenyltetrazolium bromide (MTT) reduction assay (Fig. 7m). Notably, transfection of 10 μM miR-214-3p significantly promoted light-activated pathological aggregation of PrP and significantly increased PrP toxicity in C2C12 myoblasts stably expressing WT PrP^C^ (Fig. 7k−m).

To ascertain the nature of light-activated pathological aggregates of PrP in skeletal muscle cells, we conducted immunogold electron microscopy. C2C12 cells stably expressing mCherry-Cry2-WT PrP^C^ were cultured until confluence was reached 85%, then transfected without (Fig. 7n) or with (Fig. 7o) 10 μM miR-214-3p for 30 min, cultured for 12 h after activation by 488-nm laser for 10 min, and labeled by gold particles conjugated with the anti-PrP antibody 8H4. Notably, the amyloid fibrils in the above cell samples were recognized by 8H4 and decorated with 10-nm gold labels, and the skeletal muscle cells transfected with 10 μM miR-214-3p produced much more amyloid fibrils than those transfected without miRNA (Fig. 7n,o).

Altogether these data demonstrate that PrP^C^ recruits miR-214-3p into phase-separated condensates in living skeletal muscle cells, which in turn promotes pathological aggregation of PrP.

## Discussion

Because PrP^C^ is an important protein in muscle regeneration^16^ and myoblast differentiation^15,18^, it has generally been thought that PrP^C^ dysfunction might be responsible for skeletal muscle cell death in patients with inclusion-body myositis, dermatomyositis, and other myopathies^19,20,22^. Accumulating pieces of evidence point to a crucial role of autophagy in myoblast differentiation^25−27^, whereas impaired autophagy is observed in aged muscle satellite cells^28^. We now show that maintenance of PrP^C^ homeostasis is essential for myoblast differentiation, and PrP^C^ located in the cytoplasm strongly inhibits skeletal muscle cell autophagy and blocks myoblast differentiation. Interestingly, PrP^C^ controls the distribution of caveolin 1 between lipid raft domains on the cell membrane and the cytoplasm where caveolin 1 can function to impair the ATG12-ATG5 complex and thus inhibit autophagy progression^77^.

Several in vitro studies have described physical interactions between PrP and different classes of molecules, among which nucleic acids are highlighted as potential PrP molecular partners^47,50,52,53,55,56,78,79^. Given that miRNAs play important roles in regulating differentiation, atrophy, and regeneration of skeletal muscle via interaction with specific proteins^35−38^, we predicted that PrP^C^ might physically interact with miRNAs to modulate myoblast differentiation. In this study, we observed the co-localization of overexpressed PrP^C^ with miR-214-3p in the skeletal muscle of patients with dermatomyositis, neurogenic myopathy, and muscular dystrophy. Additionally, our results show that PrP^C^ is overexpressed in skeletal muscle cells under pathological conditions and inhibits myoblast differentiation via selectively interacting with a subset of miRNAs including miR-214-3p and miR-204-5p to significantly inhibit ATG5-dependent autophagy. Therefore, PrP^C^ regulates autophagy and differentiation of skeletal muscle cells via multiple mechanisms, the first being physically interacting with specific miRNAs.

In this work, we report that PrP^C^, a glycoprotein existing in cytoplasmic form during myoblast differentiation, exhibits disparate propensities to phase separate with miRNA. We show that PrP^C^ undergoes LLPS in vitro and in cells. PrP^C^ condensates selectively recruit a subset of miRNAs such as miR-214-3p and miR-204-5p in vitro and in skeletal muscle cells, which in turn dramatically promotes the LLPS of PrP^C^ under both conditions. PrP^C^ concentrates miR-214-3p into phase-separated condensates (puncta) in the cytosol of skeletal muscle cells, which in turn mediates enhanced PrP^C^ condensation, rendering the resulting liquid droplets fibril-like. Mutations of the GCAG sequences in miR-214-3p that block the specific interaction of PrP^C^ with miR-214-3p impair the incorporation of miR-214-3p into PrP^C^ condensates formed in skeletal muscle cells, and the addition of miR-214-3p mutant only mildly enhances the phase separation and droplet fusion ability of PrP^C^. Overall, our results show that PrP^C^, a potential RNA-binding protein, undergoes miRNA-mediated LLPS in vitro and in cells. Intriguingly, RNA-binding protein YBX1 undergoes LLPS in vitro and in cells, and YBX1 condensates selectively recruit miR-223 in vitro and into exosomes secreted by cultured cells^80^. In vitro LLPS of PrP^C^ is modulated by three types of RNA molecules (polyU RNA, crude tRNA, and yeast total RNA)^52,55,56^. It should be mentioned that self-complementary RNA structures also play a role in LLPS by imparting an “identity” to biomolecular condensates. This identity prevents the merging of biomolecular condensates containing different or dissimilar RNAs^81^. Thus, the specific properties of miRNAs included in PrP^C^ condensates influence the liquidness and organization of PrP^C^ condensates.

In summary, our results describe a model to underpin molecular hypotheses of how excess PrP^C^ inhibits muscle cell differentiation via miRNA-enhanced LLPS of PrP^C^ implicated in myopathy (Fig. 8). Importantly, we show that under pathological conditions, PrP^C^, an essential protein for myoblast differentiation, is overexpressed in muscle satellite cells and myoblasts and inhibits muscle cell differentiation via selectively interacting with a subset of miRNAs, miR-214-3p and miR-204-5p, to significantly inhibit ATG5-dependent and LC3B-dependent autophagy, respectively. Then PrP^C^ selectively recruits these miRNAs into phase-separated condensates in living myoblasts, which in turn greatly enhances PrP^C^ liquid phase condensation and the subsequent solid phase condensation to produce PrP fibrils, resulting in cell death in skeletal muscle cells and the subsequent muscle bundle formation in myopathy patients characterized by incomplete muscle regeneration (Fig. 8). Under physiological conditions, when myotubes are damaged, muscle satellite cells differentiate into myoblasts, PrP^C^ is expressed in myoblasts and enhances muscle cell differentiation via selectively interacting with miR-486a-5p and miR-181b-5 to significantly promote autophagy, and finally muscle bundles are assembled to create the whole muscle (Fig. 8). The selective interaction of miRNAs with PrP^C^ during cell differentiation will be valuable in regard to understanding the functional basis underlying LLPS of proteins and inspiring future research on protein condensation diseases caused by abnormal liquid- or solid-like states of proteins^82^ and regulated by RNA^43−46^.

**Fig. 8.**
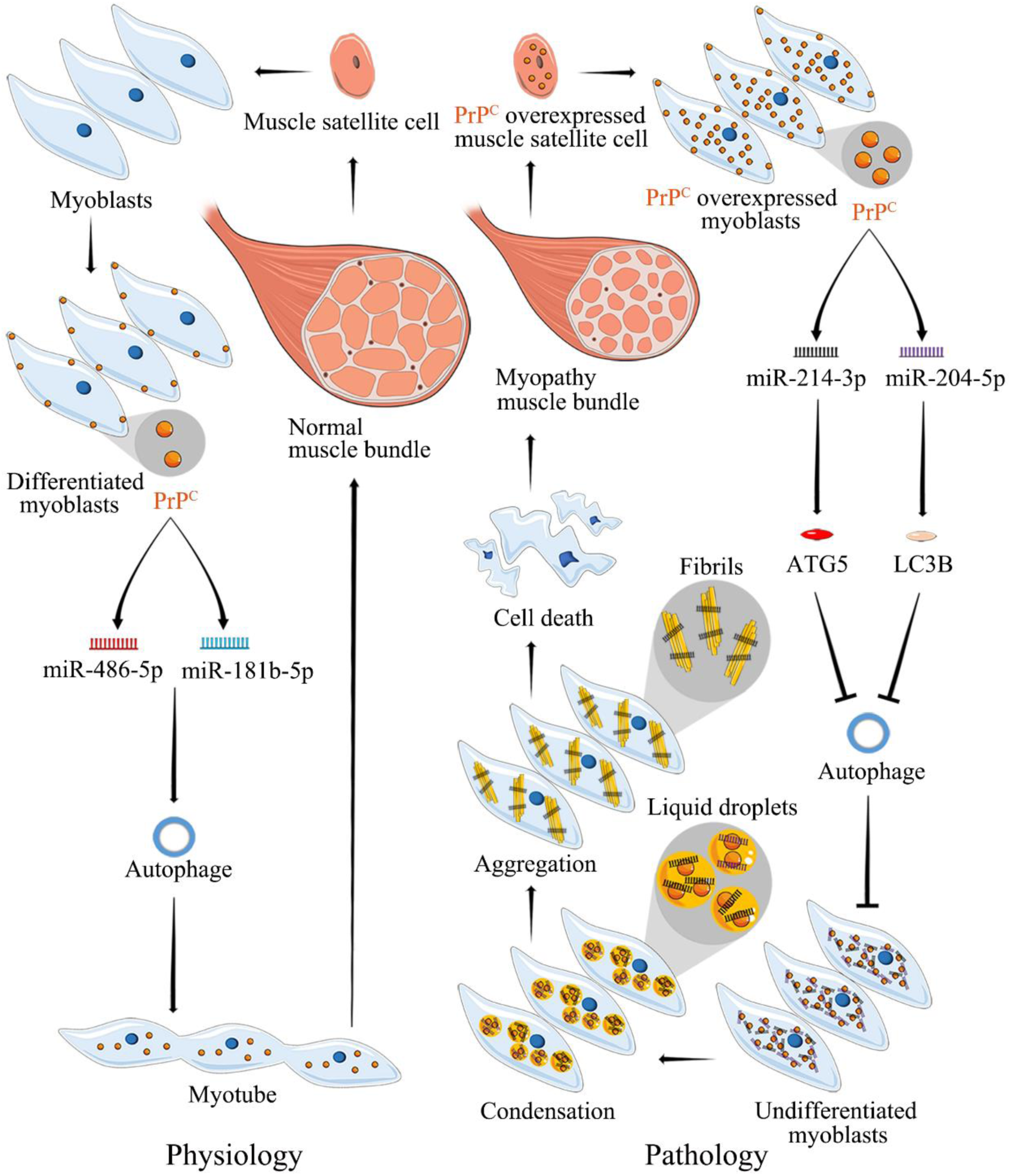
A hypothetical model shows how excess PrP^C^ inhibits muscle cell differentiation via miRNA-enhanced LLPS of PrP^C^ implicated in myopathy. Under pathological conditions (right), PrP^C^ (orange), an essential protein for myoblast differentiation, is overexpressed in muscle satellite cells (orange) and myoblasts (pale blue) and inhibits muscle cell differentiation via selectively interacting with a subset of miRNAs, miR-214-3p and miR-204-5p, to significantly inhibit ATG5-dependent and LC3B-dependent autophagy, respectively. Then PrP^C^ selectively recruits these miRNAs into phase-separated condensates (gold balls) in living myoblasts, which in turn greatly enhances PrP^C^ liquid phase condensation and the subsequent solid phase condensation to produce PrP fibrils (gold bars), resulting in cell death in skeletal muscle cells and the subsequent muscle bundle formation in myopathy patients characterized by incomplete muscle regeneration. Under physiological conditions (left), when myotubes are damaged, muscle satellite cells (orange) differentiate into myoblasts (pale blue), PrP^C^ (orange) is expressed in myoblasts (pale blue) and enhances muscle cell differentiation via selectively interacting with miR-486a-5p and miR-181b-5 to significantly promote autophagy, and finally muscle bundles are assembled to create the whole muscle.

## METHODS

### Pathological samples miRNA FISH and immunocytochemistry

The skeletal muscles of patients with myopathy such as dermatomyositis, neurogenic myopathy, and muscular dystrophy were collected at the Department of Neurology, Renmin Hospital of Wuhan University. The skeletal muscle samples were fixed with isopentane for 2-3 min, then frozen in liquid nitrogen and sliced with a cryostat sectioning. Frozen skeletal muscle sections from six myopathy patients and four age-matched controls (Table 1) were marked with hydrophobic circles using an immunohistochemical pen. H&E staining of the frozen skeletal muscle sections was conducted by following the manufacturer’s instructions (C0105S, Beyotime). For small-RNA FISH, slices were permeabilized with cold 0.5% Triton X-100 in PBS at room temperature for 5 min. After being twice washed with PBS for 5 min, the fixed cells were incubated with the prehybridization buffer for 1 h at 55 °C. Prehybridized coverslips were incubated with a hybridization buffer (10% dextran sulfate in prehybridization buffer plus 10 ng/μl FAM-labeled miRNA probe) and covered with siliconized coverslips in a humidified chamber overnight at 55 °C. Coverslips were washed with buffer (2 × SSC, 30% formamide) for 10 min at 37 °C and then with 2 × SSC, 1 × SSC, 0.5 × SSC and 1 × PBS. Immunocytochemistry staining was then performed to achieve the purpose of double staining.

For immunostaining, slices blocked with 10% goat serum for 1 h at 37 °C. Primary antibody diluted in PBS containing 3% goat serum was applied to the slices and incubated in a humidified chamber overnight at 4 °C. After three-times washes with PBS for 5 min, fluorescence-conjugated secondary antibody was applied to the cover slip and incubated in a dark room for 1 h. DAPI was then applied at the proper dilution; after being washed 3 times with PBS for 5 min and mounted with antifade mounting medium (Beyotime), cells were subjected to a Leica TCS SP8 laser scanning confocal microscope (Wetzlar, Germany). The following primary antibodies were used: mouse anti-PrP monoclonal antibody 8H4 (Abcam ab61409, 1:200) and Alexa Fluor 555-labeled donkey anti-mouse IgG (H+L) (Beyotime A0460, 1:500).

### Ethics statement

The study complies with all relevant ethical regulations. The study is based on analyses of skeletal muscle samples from two patients with dermatomyositis, two patients with neurogenic myopathy, and two patients with muscular dystrophy, and from one healthy individual, two patients with lipid storage myopathy, and one patient with glycogen storage disease (controls). The patient characteristics are described in Table 1. Tissue materials were collected at the Department of Neurology, Renmin Hospital of Wuhan University after obtaining informed consent from the patients or their relatives, who did not receive any compensation. All relevant regulations and legal requirements, including ethical approval from relevant authorities at Wuhan University, were observed during material collection.

### Cell culture and myogenic differentiation

Murine-derived C2C12 myoblast cells were cultured in minimum essential media and in Dulbecco’s modified Eagle’s medium (Gibco, Invitrogen), supplemented with 20% (v/v) fetal bovine serum (Gibco) and 1% penicillin-streptomycin in 5% CO_2_ at 37 °C. For myoblast differentiation, C2C12 myoblasts at 90% confluency were switched to a differentiation medium, supplemented with 2% horse serum (Gibco) and 1% penicillin-streptomycin.

### Plasmids and transfection

Total RNA was extracted from C2C12 myoblasts using TRIzol reagent from Beyotime (Nantong, China) according to the instructions, and the total RNA was reverse transcribed into a cDNA library using First Strand cDNA Synthesis Kit (Beyotime). The open reading frame of mouse PrP was obtained by amplifying the cDNA library using PCR, and cloned into the pBABE vector (pBABE-PrP^C^) and the pET-28a (Pet-28a-PrP). Genetic prion disease–related mutation F198S (pBABE-F198S) was constructed by site-directed mutagenesis using pBABE-PrP^C^ template. The guide RNAs (gRNAs) for mouse PrP were selected from the mouse GeCKO CRISPR library and cloned into pLenti-CRISPR-V2 vector (pLenti-CRISPR-V2-PrP) according to the instruction described^83^. The gRNA oligo and PCR primers were listed in Supplementary Table 2. The 3’UTRs of *ATG5* and *LC3B* were obtained by PCR amplification of cDNA and cloned into psiCHECK2 vector (luciferase reporter).

### Western blotting

For analysis by Western blotting and IP, C2C12 cells grown in a 6-well plate were washed twice with ice-cold PBS and lysed in 300 μl (per well) cell lysis buffer containing 1 × protease inhibitor cocktail (Target Mol). The amount of loaded protein was normalized using a BCA Protein Quantification kit (Beyotime). The cell lysates were boiled in SDS-PAGE loading buffer for 10 min and then subjected to SDS-PAGE and probed with the following specific antibodies: mouse anti-PrP monoclonal antibody 8H4 (Abcam ab61409, 1:5,000), mouse anti-MyHC antibody MF-20 (Developmental Studies Hybridoma Bank MAB4470-SP, 1:1,000), mouse anti-MyoG antibody F5D (Santa Cruz Biotechnology sc-12732, 1:500), mouse anti-β-actin (Beyotime AA128, 1:1,000), rabbit anti-ATG5 antibody (Sigma SAB5700062, 1:1,000), rabbit anti-LC3B antibody (Sigma SAB1306269, 1:1,000), Alexa-conjugated fluorescent secondary antibodies (Beyotime, rabbit A0208, 1:1,000; mouse A0216, 1:1,000).

### Cell immunocytochemistry and miRNA FISH

For immunostaining, C2C12 cells were seeded in 12-well plates, washed twice with PBS for 5 min, fixed with 4% paraformaldehyde for 30 min at room temperature, washed twice with PBS for 5 min and then permeabilized for 4 min on ice with 0.25% Triton X-100 in PBS. Cells were washed three times with PBS for 5 min, and blocked with 3% bovine serum albumin (BSA) for 30 min at 37 °C. Primary antibody diluted in PBS containing 3% BSA was applied to the cells and incubated for 3 h at 37 °C. After three-times washes with PBS for 5 min, fluorescence-conjugated secondary antibody was applied to the cover slip and incubated in a dark room for 45 min. DAPI (Beyotime) was then applied at the proper dilution; after being washed 3 times with PBS for 5 min and mounted with antifade mounting medium (Beyotime), cells were subjected to a Leica TCS SP8 laser scanning confocal microscope (Wetzlar, Germany). The following primary antibodies were used: mouse anti-PrP monoclonal antibody 8H4 (Abcam ab61409, 1:200), mouse anti-MyHC antibody MF-20 (Developmental Studies Hybridoma Bank MAB4470-SP, 1:200), rabbit anti-ATG5 antibody (Sigma SAB5700062, 1:200), rabbit anti-LC3B antibody (Sigma SAB1306269, 1:200), Alexa Fluor 488-labeled goat anti-rabbit IgG (H+L) (Beyotime A0423, 1:500), and Alexa Fluor 555-labeled donkey anti-mouse IgG (H+L) (Beyotime A0460, 1:500).

Small-RNA FISH was performed as described previously^84^ with minor modifications. C2C12 cells were seeded onto a sterile coverslip positioned in the bottom of a well in a 12-well dish and differentiated with 2% horse serum. Cells rinsed twice with PBS for 5 min, fixed in 4% paraformaldehyde in PBS for 30 min at room temperature, rinsed twice in PBS for 5 min and then permeabilized with cold 0.5% Triton X-100 in PBS at room temperature for 5 min. After being twice washed with PBS for 5 min, the fixed cells were incubated with a prehybridization buffer (2 × SSC, 1 × Denhardt’s solution, 50% formamide, 10 mM EDTA, 100 μg/ml yeast tRNA, 0.01% Tween 20, and 2 U/μl RNase inhibitor, Beyotime) for 1 h at 37 °C. Prehybridized coverslips were incubated with a hybridization buffer (10% dextran sulfate in prehybridization buffer plus 40 ng/μl FAM-labeled miRNA probe) in a humidified chamber overnight at 37 °C. Coverslips were washed with 2 × SSC, 1 × SSC, 0.5 × SSC and 1 × PBS for 5 min. Immunocytochemistry staining was then performed to achieve the purpose of double staining as follows. C2C12 cells (control) and C2C12 cells stably expressing full-length wild-type mouse PrP^C^ upon differentiation for 4 days, in which miRNA was detected by FISH (green) using FAM-labeled miRNA probe, were immunostained with the anti-PrP antibody 8H4 (red) and stained with DAPI (blue). Images of FAM-labeled miRNA (green) and PrP^C^ (red) were captured using a Leica TCS SP8 laser scanning confocal microscope (Wetzlar, Germany).

### Immunoprecipitation (IP)

Primary antibodies or IgG were added into Protein A+G Agarose beads (Fast Flow for IP) (Beyotime) on a rotator for 3 h at 4 °C. C2C12 Cells were lysed in cell lysis buffer containing 1 × protease inhibitor cocktail (Target Mol) for Western and IP, and the lysates were centrifuged at 12,000 *g* for 10 min at 4 °C to remove cell debris. An input of collected supernatant was set aside for subsequent analysis. Cell lysate was added into the mixture on a rotator overnight at 4 °C. Finally, immunocomplexes eluted from beads were detected using indicated antibodies by Western blotting.

### RNA immunoprecipitation (RIP)

C2C12 cells cultured in two 150-mm plates were washed twice with cold PBS and then crosslinking with 0.1% formaldehyde for 10 min at room temperature. Cells were added into 1.25 M glycine solution for 5 min and then washed 3 times with PBS for 5 min. Scrapped cells were harvested by centrifugation at 500 *g* for 5 min at 4 °C. Crosslinked cells were lysed in 1 ml of RIP lysis buffer containing 25 mM Tris-HCl (pH 7.4), 150 mM NaCl, 1% NP-40, 1 mM EDTA, 5% glycerin, 400 U/ml RNasin ribonuclease inhibitor, 1 mM dithiothreitol (DTT), and 1 × protease inhibitor cocktail (Target Mol) on ice for 20 min. Lysates were centrifuged at 12,000 *g* at 4 °C for 10 min to remove cell debris. An input of collected supernatant was set aside for subsequent analysis. Each sample was then divided and incubated with 15 μg of either mouse anti-PrP monoclonal antibody 8H4 or mouse IgG on a rotator for 5 h at 4 °C. Complexes were pulled down by incubation with Protein A+G Agarose beads (Fast Flow for IP) (Beyotime) on a rotator for 3 h at 4 °C. Beads were washed twice with lysis buffer and washed twice with RIP wash buffer containing 350 mM NaCl. 150 μl of proteinase K buffer containing 1.2 mg/ml proteinase K, 10 mM Tris-HCl (pH 8.0), 5 mM EDTA, 0.5% SDS, 400 U/ml RNasin ribonuclease inhibitor, and 1 mM DTT for 5 min at 55 °C to reverse formaldehyde crosslinking. Co-precipitated RNAs were extracted from beads and used for construction of small-RNA libraries or analysis by RT−qPCR using specific primers.

### RIP-seq data analysis

Reads of small RNA-seq were trimmed using the Cutadapt program and then mapped to the mouse genome (mm10) using Bowtie v0.9.6 (ref. ^85^) with parameters ‘-n 0 -l 20 -k 1 −best’. Read counts of mature miRNAs were calculated using the featureCounts program^86^ with parameters ‘–fracOverlap 0.8– fracOverlapFeature 0.8 -s 1 -M’. The enrichment of miRNAs from RIP over input and pellet over total lysates were calculated with the edgeR package^87^. We defined the enriched miRNAs as CPM > 50, log_2_ fold change > 0.5 and *P* < 0.05.

### RT−qPCR of miRNAs

The Qiagen miScript II RT Kit was used to quantify miRNAs. After TRIzol (Beyotime) extraction of total RNA, mature miRNAs were polyadenylated by PolyA polymerase and reverse transcribed into cDNAs by using an oligo-dT primer provided in the kit. The oligo-dT primer contained a 3 degenerate anchor and a universal tag sequence at the 5’ end, allowing quantitative analysis of mature miRNA by real-time PCR using the universal primer and a miRNA-specific primer. Quantitative PCR was carried out with a 1:10 dilution of cDNA, 2 × SYBR Green PCR Mix, and 10 × miScript universal primers included in the kit in combination with 10 × miRNA-specific primers (listed in Supplementary Table 3). The U6 snRNA primer from Qiagen was used for normalization, and ΔCt was calculated to derive relative expression. C2C12 cells stably expressing full-length wild-type mouse PrP^C^ and C2C12 cells KO for PrP^C^ were incubated with a differentiation medium for 4 days and then treated with 20 mg/ml α-amanitin, an inhibitor of RNA polymerase II. The above cell lines were lysed every 3 h for up to 12 h. The relative stability of miRNAs in the above cell lines was determined as the relative enrichment (to U6 snRNA) of miRNAs by RT−qPCR.

### Small-RNA pulldown

For the small-RNA pulldown assay, 5’ biotinylated miRNA baits were commercially synthesized (listed in Supplementary Table 4). For the preparation of cell lysate, a 155-mm plate of confluent C2C12 cells were harvested by centrifugation at 500 *g* for 5 min at 4 °C. Crosslinked cells were lysed in 1 ml of pulldown lysis buffer containing 25 mM Tris-HCl (pH 8.0), 150 mM NaCl, 1% NP-40, 1 mM EDTA, 5% glycerin, 400 U ml^-1^ RNasin ribonuclease inhibitors, 1 mM DTT on ice for 20 min. Insoluble material was removed by centrifugation at 20,000 *g* for 30 min at 4 °C. Lysates were centrifuged at 12,000 *g* for 10 min at 4 °Cto remove cell debris. An input of collected supernatant was set aside for subsequent analysis. For each pulldown assay, 40 μl of biotin-labeled small RNA (20 μM) was incubated with precleared lysate overnight at 4 °C with rotation. One-hundred microliters of magnetic streptavidin beads (Beyotime) were washed with pulldown lysis buffer twice and incubated with the RNA lysates for 3.5 h at 4 °C with rotation. The beads were washed with cold wash buffer I containing 300 mM NaCl, washed with cold wash buffer II containing 0.05% tween-20, washed with cold pulldown lysis buffer, and then boiled in 40 μl SDS loading buffer for analysis by Western blotting.

### Luciferase reporter assay

For luciferase assays, C2C12 cells were seeded in 12-well plates and co-transfected with 1.25 μg of luciferase reporter plus 10 μM miRNA mimics or NC. After 36 h, cells were harvested for luciferase assays using the Dual Luciferase Reporter Gene Assay Kit (Yeasen, 11402ES60). A Cytation 3 Cell Imaging Multi-Mode Reader (BioTek) was used to collect light generated by *Renilla* or firefly.

### Protein purification

The open reading frame of mouse PrP was obtained by amplifying the cDNA library using PCR, and cloned into the pBABE vector (pBABE-PrP^C^) and the pET-28a (Pet-28a-PrP). Pet-28a-PrP plasmid was transformed into *E. coli*. Recombinant full-length wild-type mouse PrP was expressed in *E. coli* BL21 (DE3) cells (Novagen, Merck, Darmstadt, Germany) and purified by high-performance liquid chromatography on a C4 reverse-phase column (Shimadzu, Kyoto, Japan) as described by Bocharova et al^88^ and Zhou et al^89^. After purification, recombinant wild-type mouse PrP^C^ was dialyzed against 1 × PBS (pH 7.4) for 24 h, concentrated, filtered, and stored at -80 °C. SDS-PAGE and mass spectrometry were used to confirm that the purified wild-type mouse PrP was single species with an intact disulfide bond. We used a NanoDrop OneC Microvolume UV-Vis Spectrophotometer (Thermo Fisher Scientific) to determine the concentration of wild-type mouse PrP^C^, using its absorbance at 280 nm and the molar extinction coefficient calculated from the composition of the protein (http://web.expasy.org/protparam/).

### Liquid-droplet formation

The freshly bacterial purified wild-type mouse PrP^C^ was incubated with TAMRA (red fluorescence, excitation at 561 nm) at a PrP^C^: TAMRA molar ratio of 1:3 for 1 h. These labeled proteins were filtered, concentrated to 245 μM in a centrifugal filter (Millipore) and diluted in 1 × PBS (pH 7.4). In total, 25, 35, 45, 50, 65, and 80 μM wild-type mouse PrP^C^ labeled by TAMRA were incubated with 1 × PBS (pH 7.4) on ice to induce LLPS^48^ for 5 min. In total, 50 μM wild-type mouse PrP^C^ labeled by TAMRA was incubated with 1 × PBS (pH 7.4) containing 0, 1.25, 2.5, 5, 10 or 20 μM miRNA on ice to induce LLPS for 5 min. The miRNAs include a subset of PrP^C^-bound miRNAs, such as miR-214-3p and miR-204-5p, and nonspecific miR-183-5p (negative control). Liquid droplets of PrP^C^ (protein condensates) formed in 1 × PBS containing 0−20 μM miRNA were observed by a Leica TCS SP8 laser scanning confocal microscope (Wetzlar, Germany) with excitation at 561 nm. In total, 50 μM wild-type mouse PrP^C^ labeled by TAMRA (red fluorescence) was incubated with 1 × PBS containing 10 μM FAM-labeled miRNA (green fluorescence) on ice to induce LLPS for 5 min. The FAM-labeled miRNAs, including FAM-labeled miR-214-3p, FAM-labeled miR-204-5p, FAM-labeled miR-83-5p, and FAM-labeled mutant miR-214-3p (listed in Supplementary Table 4), were synthesized. PrP^C^ de-mixed droplets (red) fused with droplets of FAM-labeled miRNA (green) were observed by a Leica TCS SP8 laser scanning confocal microscope (Wetzlar, Germany), with excitation at 561 nm and 488 nm, respectively. All phase separation experiments were performed at least three times and were pretty reproducible.

### Fluorescence recovery after photobleaching (FRAP)

In total, 50 μM wild-type mouse PrP^C^ labeled by TAMRA was incubated with 1 × PBS (pH 7.4) or incubated with the same buffer further containing 10 μM miRNA on ice to induce LLPS for 10 min. Liquid droplets of wild-type PrP^C^ were observed by a Leica TCS SP8 laser scanning confocal microscope with excitation at 561 nm. For each droplet, a square was bleached at 100% transmission for 320 ms, and postbleaching time-lapse images were collected (100 frames, 320 ms per frame). Images were analyzed using Zen (LSM 880 confocal microscope manufacturer’s software). All FRAP experiments were repeated three times and the results were reproducible.

### OptoIDR assays

We employed an optogenetic tool that uses a blue light (488-nm laser) to activate IDR-mediated LLPS of proteins in living C2C12 cells^71−73^. C2C12 cells were transfected with optoIDR plasmids (pBABE-mCherry-PrP^C^). At 24 h post-transfection, cells were plated at 35-mm confocal dishes. After another 24 h, images were captured using a Zeiss LSM 880 with Airyscan confocal microscopy. Unless indicated otherwise, droplet formation was induced with 488-nm light pulses every 2 s for the duration of the imaging as indicated, and images were taken every 2 s. Fluorescence from mCherry was excited with 561-nm light.

### Live-cell imaging

Cells were grown on chambered cover 35 mm confocal dishes to an appropriate density. Cells were transfected with FAM-labeled miR-214-3p using TransIT-X2 (Mirus). Live-cell images of mCherry-PrP^C^ (red)-FAM-labeled miR-214-3p (green) speck formation were captured after 30 min using a Zeiss LSM 880 with Airyscan confocal microscopy with a 63 × oil objective and with excitation at 561 nm and 488 nm, respectively. Images were analyzed using Zen (LSM 880 confocal microscope manufacturer’s software).

### Cellular FRAPAssays

Cells were grown on chambered cover 35 mm confocal dishes for 24 h, transfected with 10 μM miR-214-3p using TransIT-X2 (Mirus) for 30 min in a growing medium, and subjected to laser excitation for 10 s. FRAP was then done on a Zeiss LSM 880 with Airyscan confocal microscopy with a 561 nm laser. Bleaching was undertaken over a ∼1 mm radius using 80% laser power, and images were collected every 5 s. Fluorescence intensity was measured using ZEN (LSM 880 confocal microscope manufacturer’s software). All FRAP experiments were repeated three times and the results were reproducible.

### Sarkosyl-insoluble Western blotting

Sixty percent of the supernatant was incubated with 1% sarkosyl for 30 min at 25 °C. The mixture was then ultracentrifuged at 150,000 *g* for 30 min, and the supernatant was carefully removed. The sarkosyl-insoluble pellets were boiled in the SDS-PAGE loading buffer for 10 min. The forty percent of the supernatant, which served as the total protein sample, was also boiled in the SDS-PAGE loading buffer for 10 min. The samples were separated by 12.5% SDS-PAGE and then Western blotted as follows. The samples were transferred to polyvinylidene difluoride membranes (Millipore). The membranes were blocked with 5% fat-free milk in 25 mM Tris-buffered saline buffer containing 0.047% Tween 20 (TBST). Then the sarkosyl-insoluble pellets from the above cells were probed by the anti-PrP antibody 8H4, and the corresponding cell lysates were probed using 8H4 and anti-β-actin antibody, respectively. The amount of loaded protein was normalized using a BCA Protein Quantification kit (Beyotime). For calculating the amounts of sarkosyl-insoluble PrP, the ImageJ software (NIH) was used to assess the densitometry of PrP bands. The normalized amount of insoluble PrP aggregates in C2C12 cells stably expressing mCherry-Cry2-wild-type PrP^C^ was determined as a ratio of the density of insoluble PrP aggregate bands over that of the total PrP bands in cell lysates.

### Cell viability assays

C2C12 cells were plated in 96-well plates in minimum essential medium. The MTT stock solution (5 mg/ml) was diluted with 1 × PBS and added into the well for 4 h until formazan was formed in cells. The final concentration of MTT was 0.5 mg/ml. Finally, the dark blue formazan crystal was dissolved with dimethyl sulfoxide, followed by measuring its absorbance at 492 nm using a Thermo Multiskan MK3 microplate reader (Thermo Fisher Scientific). Cell viability was expressed as the percentage ratio of the absorbance of wells containing the samples treated with miRNA to that of wells containing cells treated without miRNA.

### Immunogold electron microscopy of PrP fibrils

The cells were then resuspended with 450 μl of 10% sucrose solution in PBS (pH 7.4) containing 1 × protease inhibitor cocktail (Target Mol). The mixtures were sonicated 10 times on ice at 50 W and 3 s/3 s, and then centrifuged at 17,000 *g* for 30 min at 4 °C to remove the cell debris. Sixty percent of the supernatant was incubated with 1% sarkosyl for 30 min at 25 °C. The mixture was then ultracentrifuged at 150,000 *g* for 30 min, and the supernatant was carefully removed. The sarkosyl-insoluble pellets were resuspended in PBS (50 μl). Sample aliquots of 10 μl were absorbed onto nickel grids for 1 min, and then wash twice with water for 30 s. After blocked with 0.1% BSA of 10 μl, samples on grids were incubated with 1/100 anti-PrP antibody 8H4 of 10 μl. After wash twice with water for 30 s, 1/200 10-nm gold-labeled homologous secondary antibodies of 10 μl were used to incubate the grids for 20 min at 37 °C. Unbound gold-labeled homologous secondary antibodies were removed by washing with 200 μl of water drop by drop. Samples on grids were then stained with 2% (w/v) uranyl acetate for 1 min. The stained samples were examined using a JEM-1400 Plus transmission electron microscope (JEOL) operating at 100 kV.

### Statistical analysis

The data shown for each experiment were based on at least three technical replicates, as indicated in individual figure legends. Data are presented as mean ± S.D., and *p-*values were determined using a two-tailed unpaired *t* test in Figs. 4a,b and 5e. One-way ANOVA and multiple comparisons were performed by SPSS 19.0 and different letters indicate significant differences at the level of *p* < 0.05 in Figs. 2d,e, 4d−h. All experiments were further confirmed by biological repeats.

## Reporting summary

Further information on experimental design is available in the Nature Research Reporting Summary linked to this article.

## Data availability

All RIP-seq data from this study have been deposited in the Gene Expression Omnibus (GEO) under series accession number GSE203419. The source data including the statistical source data for Fig. 3d,e, Fig. 5a,b,d,e,g−k, Fig. 6d, and Fig. 7l,m and the uncropped gel images for Fig. 2a−d, Fig. 3c, Fig. 4b,j, Fig. 5c,f, and Fig. 7k are provided with this paper. Other data are available upon reasonable request.

## Acknowledgements

Y.L. acknowledges fundings from the National Natural Science Foundation of China (nos. 32271326, 32071212, and 31770833). Y.L. also acknowledges financial support from the Key Project of Basic Research, Science and Technology R&D Fund of Shenzhen (no. JCYJ20200109144418639) and the Translational Medicine and Interdisciplinary Research Joint Fund of Zhongnan Hospital of Wuhan University (no. ZNJC201934). L.X. was supported by the National Natural Science Foundation of China (nos. 81900797 and 81871036) and Guangdong Basic and Applied Basic Research Foundation (no. 2020B1515020046). Y.Z. acknowledges financial support from the Scientific Research Project of Hubei Health Commission (no. WJ2023M079). L.-Q.W. acknowledges financial support from the National Natural Science Foundation of China (no. 32201040), China Postdoctoral Science Foundation (nos. 2021TQ0252 and 2021M700103), and the Fundamental Research Funds for the Central Universities (no. 2042022kf1047). Y. Zhou acknowledges financial support from the National Natural Science Foundation of China (no. 31922039) and the Natural Science Foundation of Hubei Province (no. 2020CFA057). We thank X. Ji (Peking University) for the gift of the pHR-mCherry-cry2 vector and Y. Liu (Wuhan University) for helpful suggestions.

## Author contributions

Y.L. supervised the project. J.T. and Y.L. designed the experiments. J.T., L.-Q.W. and J.C. purified the mouse PrP^C^ and cultured the cells. J.T. and B.D. performed in vitro phase separation and droplet fusion experiments. J.T. performed Western blotting and immunoblotting analyses, RIP-seq, small RNA-seq, and OptoIDR assays. J.T., Y. Zhou and Y.L. performed RIP-seq and small RNA-seq analyses. Y.Z., Y. Liu and Z.L. provided the skeletal muscle samples. Y.Z. and Y. Liu performed H&E staining of frozen skeletal muscle sections. X.P. and L.X. provided transfection methods and plasmids for myoblasts. J.T. and Y.L. wrote the manuscript. All authors proofread and approved the manuscript.

## Competing interests

The authors declare no competing interests.

---

**Extended Data Fig. 1.**
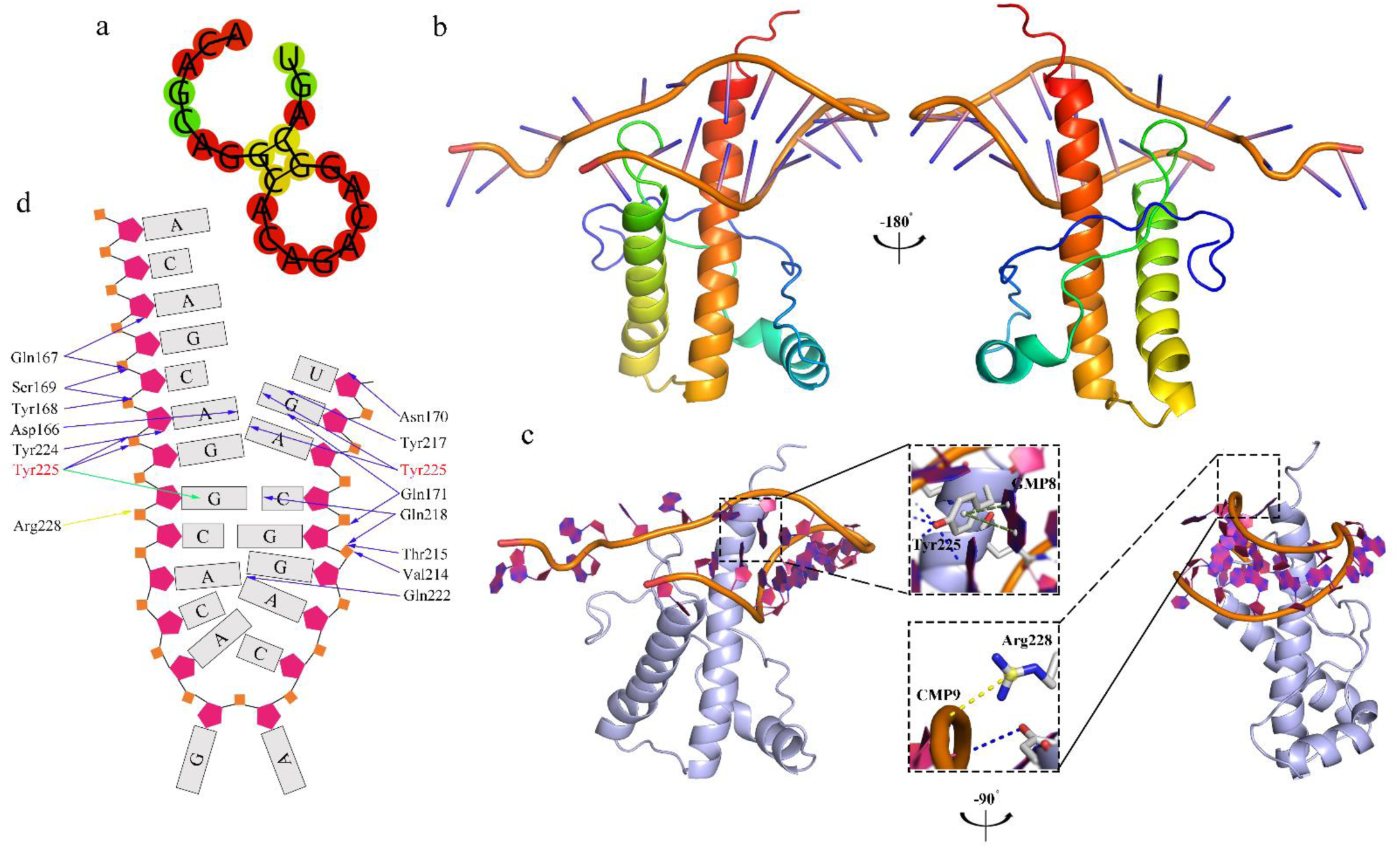
Prediction of the binding sites of miR-214-3p for PrP^C^. **a**, Secondary structure of miR-214-3p with a key region highlighted in yellow (stem) predicted by RNAComposer. **b**,**c**, Cartoon representation of the structure of the PrP^C^:miR-214-3p complex in three different views. Molecular docking of PrP^C^ with miR-214-3p was performed using HDOCK, a protein–protein/nucleic acid protein docking web server by combining template-based and free docking. Ribbon representation of the structure of mouse PrP^C^ showing three α-helices (α1, α2, and α3) in the C-terminal domain of PrP^C^ (PDB 1XYX)^68^. **c**, Predicting the positions of salt bridges, π-bonds, and hydrogen bonds using PyMOL. Two magnified side views (middle) of two regions of the interface between PrP^C^ and miR-214-3p highlighting two π-bonds between Tyr225 in PrP^C^ and GMP8 in miR-214-3p (green) and a salt bridge between Arg228 in PrP^C^ and CMP9 in miR-214-3p (yellow). Blue dashed lines represent abundant hydrogen bonds formed in the two regions of the interface. **d**, Interactions between PrP^C^ and miR-214-3p. Asp166, Gln167, Tyr168, Ser169, Asn170, Gln171, Val214, Thr 215, Tyr217, Gln218, Gln222, Tyr224, and Tyr225 in PrP^C^ form abundant hydrogen bonds with two GCAG sequences present in miR-214-3p, which contribute to maintenance of the structure of the PrP^C^-miR-214-3p complex. Tyr225 in PrP^C^ forms π-bonds with GMP8 in miR-214-3p, and Arg228 in PrP^C^ forms a salt bridge with CMP9 in miR-214-3p. Therefore, two GCAG sequences present in miR-214-3p are the predicted binding sites for PrP^C^. Yellow, green, and blue arrows represent the observed salt bridge, π-bond, and hydrogen bonds formed in the complex, respectively.

**Extended Data Fig. 2.**
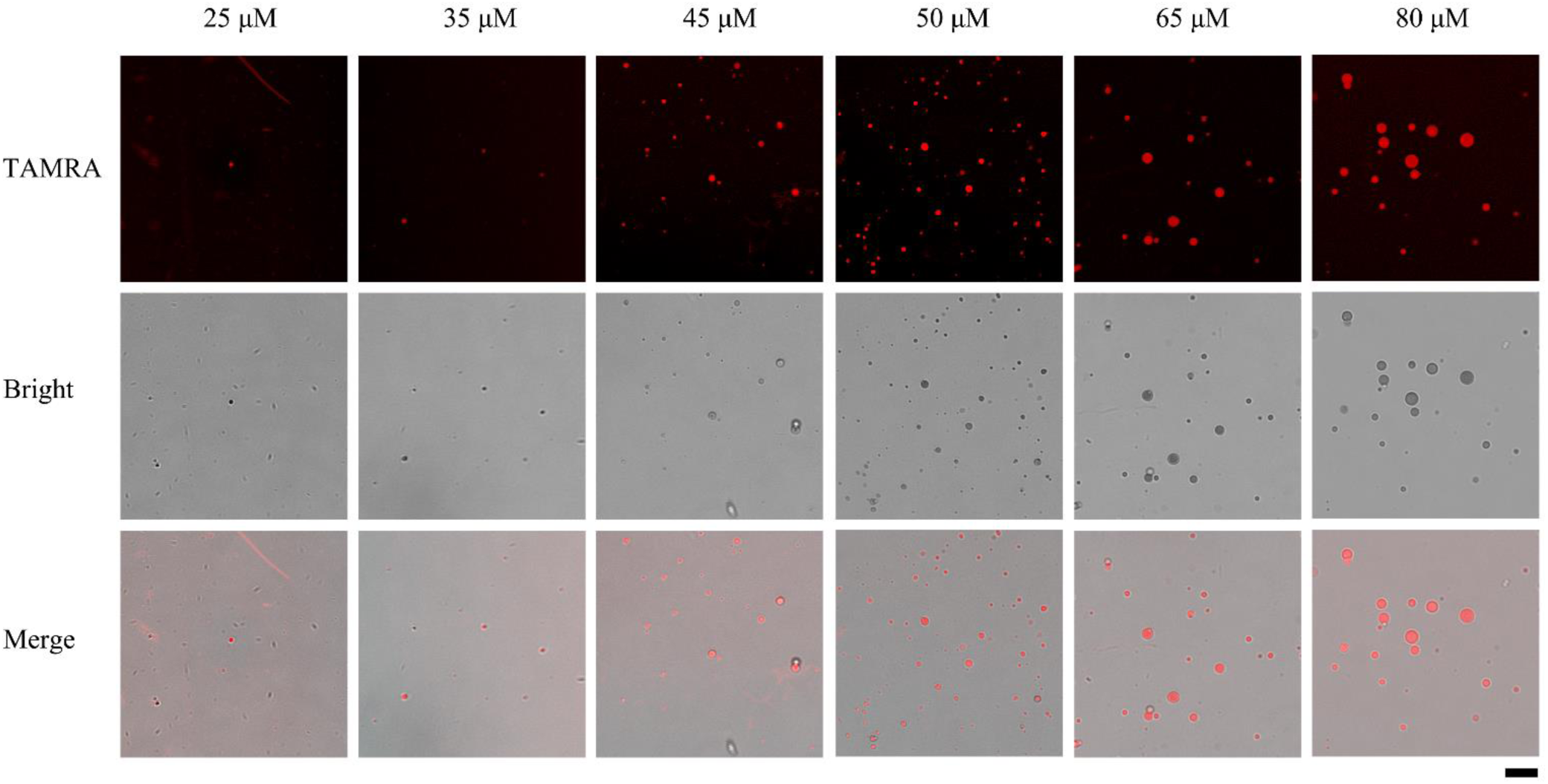
PrP^C^ undergoes liquid–liquid phase separation in vitro and forms protein condensates. Samples (25, 35, 45, 50, 65, and 80 μM) of bacterial purified wild-type mouse PrP^C^ were labeled by TAMRA (red fluorescence) and incubated with 1 × PBS (pH 7.4) on ice to induce LLPS for 5 min. PrP^C^ de-mixed droplets (protein condensates) were observed by confocal microscopy, with excitation at 561 nm. The second and third rows show the brightfield images and the merged images for PrP^C^ LLPS, respectively. Scale bar, 7.5 nm.

## Notes

### Competing Interest Statement

The authors have declared no competing interest.

